# The laboratory domestication of zebrafish: from diverse populations to inbred substrains

**DOI:** 10.1101/706382

**Authors:** Jaanus Suurväli, Andrew R Whiteley, Yichen Zheng, Karim Gharbi, Maria Leptin, Thomas Wiehe

## Abstract

The zebrafish (*Danio rerio*) is a model vertebrate widely used to study disease, development and other aspects of vertebrate biology. Most of the research is performed on laboratory strains, one of which has been fully sequenced in order to derive a reference genome. It is known that the laboratory strains differ genetically from each other, but so far no genome-scale survey of variation between the laboratory and wild zebrafish populations exists.

Here we use Restriction-Associated DNA sequencing (RAD-seq) to characterize three different wild zebrafish lineages from a population genetic viewpoint, and to compare them to four common laboratory strains. For this purpose we combine new genome-wide sequence data obtained from natural samples in India, Nepal and Bangladesh with a previously published dataset. We measured nucleotide diversity, heterozygosity, allele frequency spectra and patterns of gene conversion, and find that wild fish are much more diverse than laboratory strains. Further, in wild zebrafish there is a clear signal of GC-biased gene conversion that is missing in laboratory strains. We also find that zebrafish populations in Nepal and Bangladesh are distinct from all the other strains studied, making them an attractive subject for future studies of zebrafish population genetics and molecular ecology. Finally, isolates of the same strains kept in different laboratories show a clear pattern of ongoing differentiation into genetically distinct substrains. Together, our findings broaden the basis for future genetic and evolutionary studies in *Danio rerio*.

## Introduction

Zebrafish (*Danio rerio*) is a small fish native to freshwater bodies of the subtropical India, Nepal and Bangladesh (Whiteley, et al. 2011; Parichy 2015). In laboratory conditions the zebrafish breed often, produce many offspring, are relatively easy to grow and have translucent embryos, making them an excellent model species for drug screening, developmental biology and genetics. A common feature of zebrafish laboratory strains is that they have gone through independent genetic bottlenecks followed by different regimes of inbreeding. Therefore, one expects to observe a substantial reduction of genetic diversity when comparing laboratory strains with wild populations. Due to their independent origins, laboratory strains are expected to genetically differ not only from the wild fish but also from each other. Hence, conclusions drawn from a single laboratory strain do not necessarily apply to the other strains, nor are they fully representative for wild populations (Brown, et al. 2012; Butler, et al. 2015; Baker, et al. 2018; Balik-Meisner, et al. 2018; Holden, et al. 2019).

Given the ample resources and results from decades of laboratory zebrafish research, e.g. a high quality reference genome based on the Tübingen (TU) strain (Howe, et al. 2013), studies of natural populations are timely and offer an opportunity to complement the existing knowledge by insights on evolutionary history, behaviour, genetic variation and adaptive potential that could be obtained from studying wild populations in their natural habitats. Genetic polymorphism, and differences among laboratory strains or wild populations, is not usually a problem in the study of highly conserved processes or genes, for instance those directing the development of an organism, or core growth or signalling pathways. The laboratory fish also remain an excellent disease model for diseases based on such pathways. Complex protocols have been established to model many diseases affecting humans, these models can also be used for drug testing in early phases of pharmaceutical development (Cornet, et al. 2018). In contrast, many of the biological functions and characteristics associated with success in natural habitats would be best studied in wild populations, as the sheltered living conditions of laboratory specimens are very distinct from those in the wild. In zebrafish facilities all embryos are sanitized, feeding is regular and standardized and pathogen contact is minimized with strict procedures and protocols (Murray, et al. 2016). The laboratory strains face a substantially reduced variety of diseases and pathogens and their immune response is therefore expected not to be under nearly as strong diversifying selection as it is in the wild. Combined with the bottleneck during the establishment of a new strain it is expected that within a given strain the immune genes would be much less polymorphic than they are in the wild, hence affecting all results from studies seeking insights into the zebrafish immune system. Furthermore, while in laboratory conditions it is possible to experimentally identify genes involved in a particular metabolic pathway and even to model metabolic diseases affecting humans (Seth, et al. 2013), association studies among wild populations have the potential to reveal and dissect even minor contributors to polygenic traits that give a selective advantage in a particular habitat (Gagnaire and Gaggiotti 2016). Another field that greatly benefits from working with wild fish directly is behavioural research (Bhat, et al. 2015; Suriyampola, et al. 2016).

A study published in 2011 examined both genomic and mitochondrial variation in several wild zebrafish populations from Southern Asia and revealed three major lineages to be present. Common laboratory strains were found to be genetically closest to fish captured from North-East India (Whiteley, et al. 2011; Wilson, et al. 2014). Sequencing the genome of one wild zebrafish from North-East India revealed nearly seven million differences from the reference genome (Patowary, et al. 2013).

Here we report the results of a whole genome survey of genetic variability in a large set of laboratory and wild zebrafish. We performed Restriction-Associated DNA Sequencing (RAD-seq) on wild-caught zebrafish from three major lineages, thus complementing an existing large dataset of RAD-seq data from laboratory and wild-derived fish. Joining both datasets we carried out a comparative analysis of wild and laboratory fish and observed major differences in genetic variability, in levels of heterozygosity and in the allele frequency spectra among, but also within, the two groups (wild and laboratory zebrafish). Mutation patterns in wild fish show a clear bias in favour of G/C which appears to be nearly absent in laboratory zebrafish strains. Among fish, this bias has been previously studied mainly in the three-spined stickleback (Capra and Pollard 2011; Roesti, et al. 2013). Finally, we present evidence that isolates of the same strains obtained from different laboratories show remarkable genetic differentiation and can thus be considered distinct substrains.

## Results

### Description of the dataset

An overview of the samples is shown on **Figure 1**. We sequenced ∼0.3% of the genome (4,374,886 positions) of 26 wild and 29 laboratory zebrafish, and together with the resulting data, re-analysed 75 wild-derived and 215 laboratory zebrafish from a previously published dataset (Wilson, et al. 2014). 241,238 SNPs (Single Nucleotide Polymorphisms) were identified (overlapping a total of 11,531 genes), of which 124,015 map to introns. 102,302 of the SNPs are transitions (Ti) and 138,936 are transversions (Tv), resulting in a Ti/Tv ratio of 1.358, a value that is higher than what can be estimated for other zebrafish datasets (1.203 reported in (Stickney, et al. 2002), 1.132 based on biallelic sites in dbSNP). Only 8% of the SNPs (19,202) were reported by *Ensembl VEP* as previously known. 222,695 (92%) of the variant alleles were present in wild populations, 146,243 of these (60% of the total dataset) were not found in any of the laboratory strains. 18,543 (8%) of the identified variant alleles were present in one or more laboratory strains yet not present in any of the wild fish.

**Figure 1.**
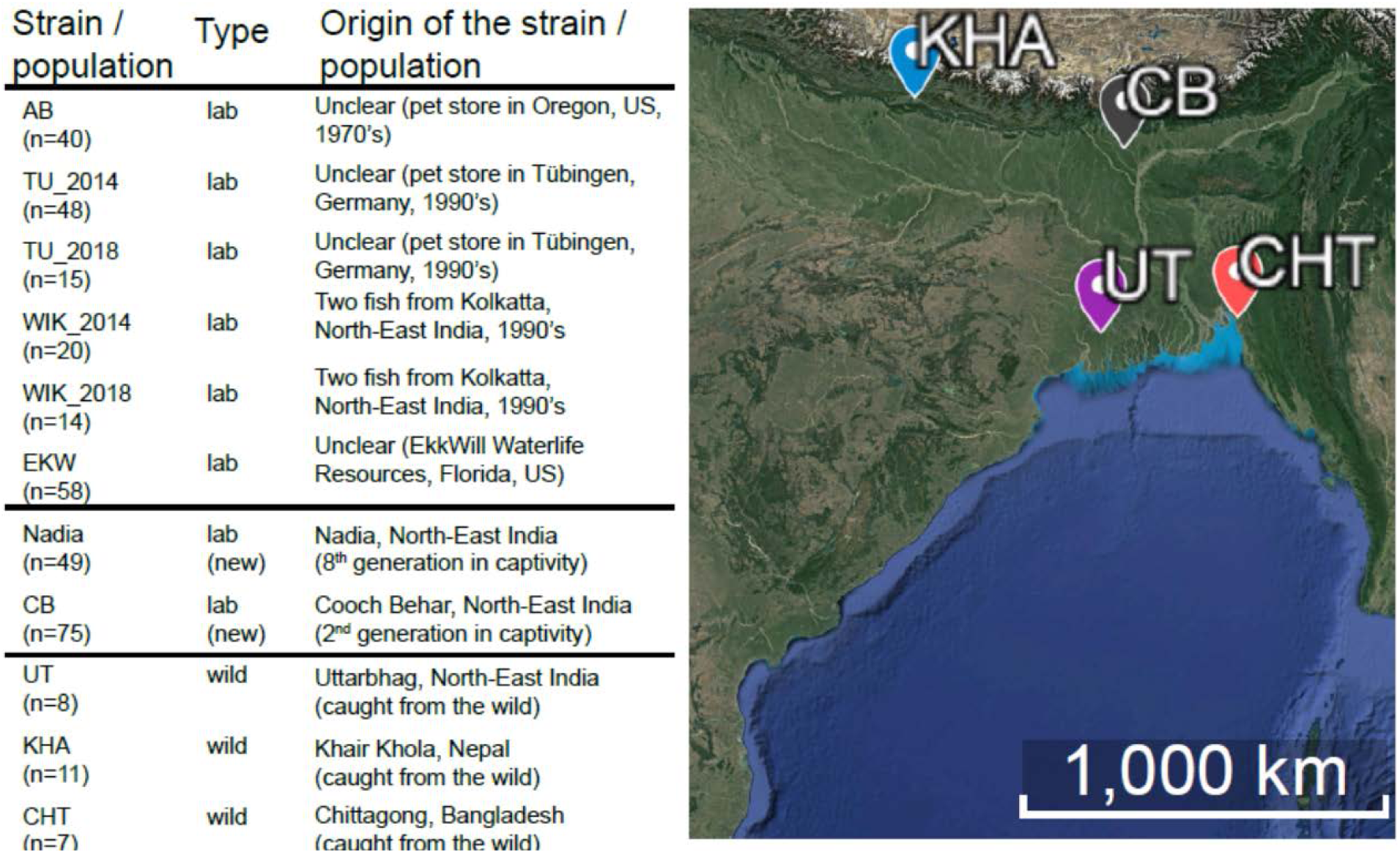
Zebrafish samples used in the study. Sample descriptions were obtained from publications first describing the fish (Whiteley, et al. 2011; Wilson, et al. 2014), when applicable. n = number of individuals sampled. The map of sampling locations was obtained from Google Earth v7.3.2 (April 04, 2019). SIO, NOAA, U.S. Navy, NGA, GEBCO. Image Landsat / Copernicus [December 14, 2015].

### Population structure of laboratory and wild zebrafish

According to how the data were collected - five laboratory strains, three wild populations and one wild-derived population (CB) - we expected that an admixture analysis would yield a likelihood peak for nine populations. However, admixture analysis revealed that the samples likely come from 13 subpopulations instead. CB appeared as a mixture of three distinct subpopulations; the independently obtained TU and WIK (University of Oregon, 2014 and Zebrafish International Resource Center, 2018) were distinct from one another as well **(Figure 2A).**

**Figure 2.**
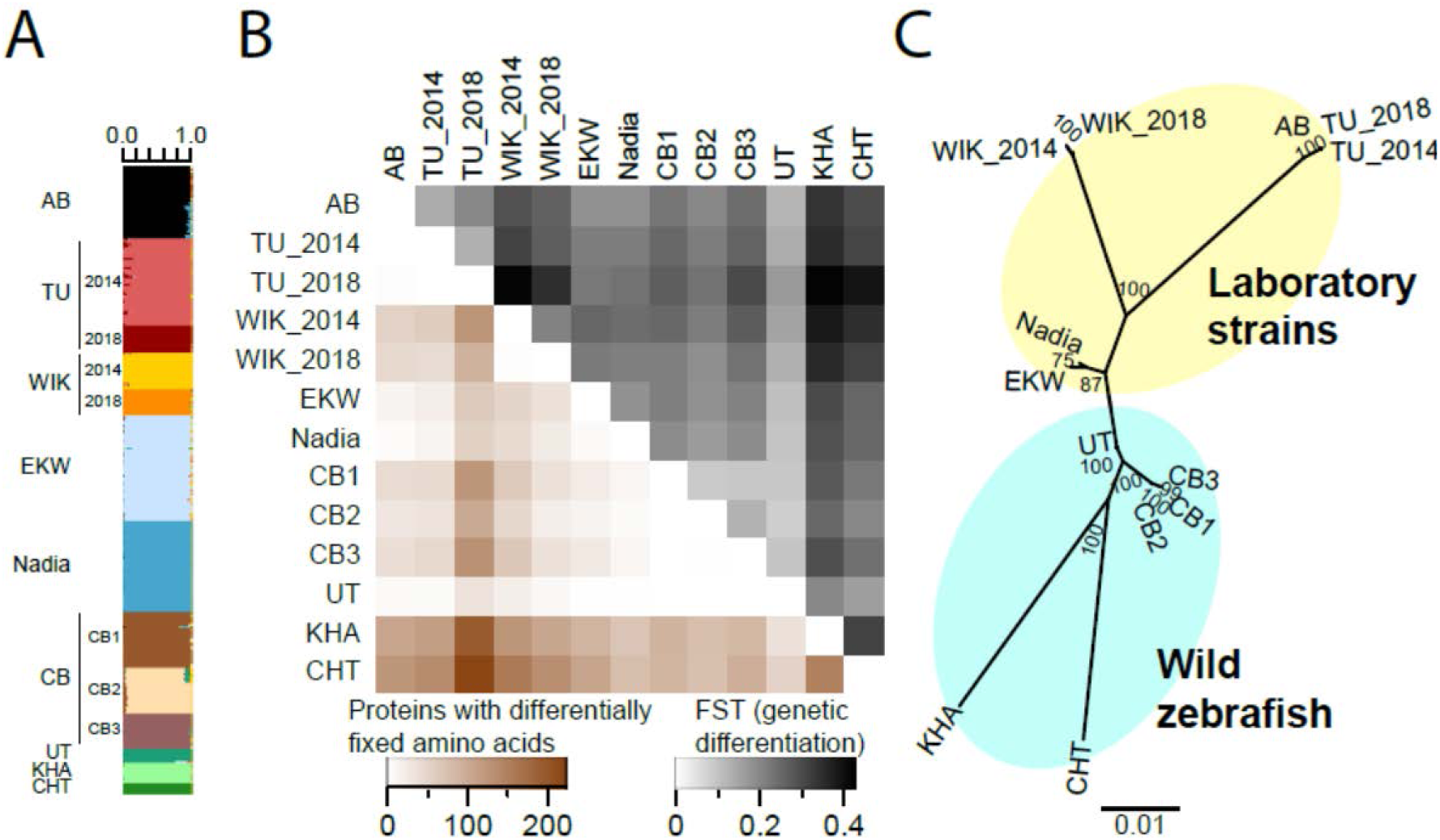
Population structure of the zebrafish samples. **(A)** Admixture plot generated by the R package LEA. Each row corresponds to one fish. Colours indicate the proportion of variation shared between individuals. 13 distinct subpopulations are identified, three of them within CB **(B)** Different zebrafish lineages have fixed differences in the amino acid sequence that are carried by every individual of the strain/population (below the diagonal). This correlates strongly with genetic distances (above the diagonal). **(C)** Unrooted Maximum Likelihood tree, generated with 1,000 bootstrap replicates.

We plotted as heatmaps different measures for mean pairwise genetic distances between populations **(Figure 2B, Supplementary Data S1).** Wild fish from India (UT) appeared genetically very close to all the laboratory strains in the study. AB was found to be closely related to both TU isolates, however, TU_2018 is much more differentiated from other strains than AB or TU_2014 **(Figure 2B)**. Among the wild fish, CHT and KHA have hundreds of fixed non-synonymous differences from the other populations (**Fig 2B, Supplementary Data S2**). Three protein-coding genes were observed to have fixed non-synonymous differences between the two WIK isolates, resulting in the amino acid changes 79Glu->Lys in chitin synthase (*chs1;* dbSNP id rs507105222), 559Glu->Gly in DNA polymerase eta (*polh*, dbSNP ID rs502489776) and 2133Leu->Gln in dystonin (*dst*, no dbSNP id for the SNP).

A phylogenetic tree constructed with these population assignments revealed a clear separation of KHA and CHT (Whiteley, et al. 2011) from the rest **(Figure 2C)**. AB and the two TU isolates appear closely related to each other; a similar pattern could be observed for the WIK isolates and for all three CB subgroups. EKW and Nadia appear be monophyletic in the tree, albeit with less bootstrap support **(Figure 2C).**

Four populations/strains had fixed nonsynonymous substitutions not present elsewhere in the data (**Supplementary Data S2**). These included 48 positions in CHT and 34 in KHA, but also one in the EKW strain (330Asp->Asn in the lipase maturation factor *lmf2a*) and another in the 2018 isolate of WIK (109Ala->Glu in *tanc2*, a gene for which the mouse knockout is embryonic lethal (Han, et al. 2010)). No sequence data were available for the locus in the 2014 isolate of WIK, likely resulting from allelic dropout (Andrews, et al. 2016).

### Within-strain genetic diversity is higher in the wild zebrafish than in the laboratory strains

At the population level, clear differences could be observed between zebrafish from the wild and the laboratory strains. We calculate three different estimators (*θ*_π_, *θ*_w_ and *θ*_1_) of the scaled mutation rate, *θ.* All had much higher values in the wild fish and in CB than in the laboratory strains, including the “newer” strain Nadia. In most cases we observe a *θ*_π_ value of around 0.003-0.007 (0.3%-0.7%), with laboratory strains at the lower end of the scale (**Figure 3A**). This is consistent with previous studies that estimated the genetic diversity in zebrafish to be at around ∼0.5%, with wild fish being more diverse than laboratory strains (Guryev, et al. 2006; Coe, et al. 2009; Whiteley, et al. 2011; Butler, et al. 2015; Balik-Meisner, et al. 2018). Other estimators of *θ* are in approximately the same range. In KHA and CHT the value of *θ*_π_ is similar to what is observed for laboratory strains. Among the laboratory strains, the TU isolate from 2018 (TU_2018) showed the least diversity – less than 0.2% for all estimators **(Figure 3A)**. In contrast, WIK_2018 is more diverse than WIK_2014.

**Figure 3.**
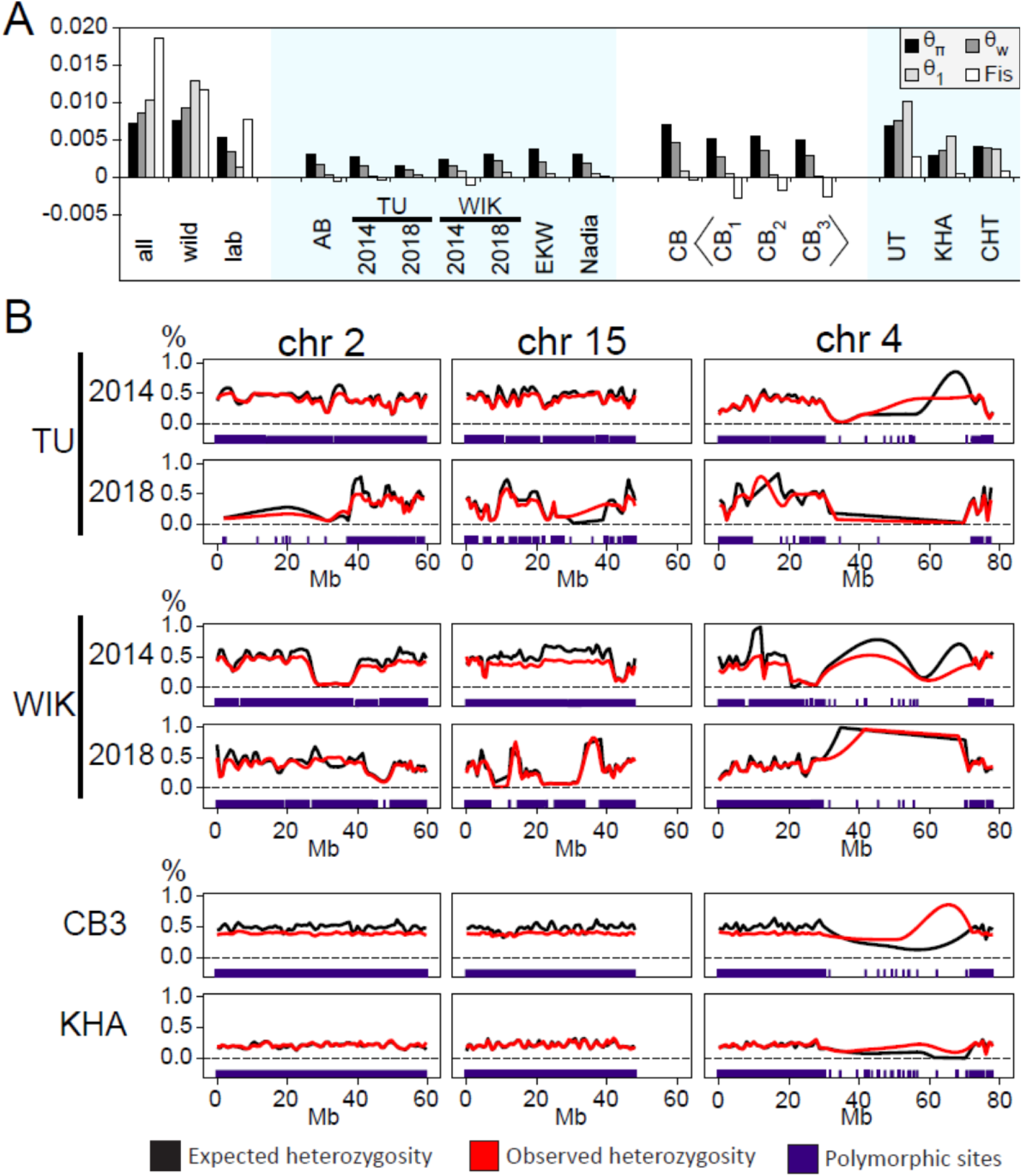
Within-strain variability of zebrafish. **(A)** Three different estimators of the scaled mutation rate, calculated independently for each population, show wild fish (CB, UT, KHA, CHT) to be genetically much more diverse than any of the laboratory strains. *θ*_π_ – average number of pairwise differences, divided by total length of the sequence. *θ*_w_ – proportion of polymorphic sites, normalized with sample size. *θ*_1_ – observed number of singleton mutations, divided by total length of the sequence. Fis – coefficieny of inbreeding. (**B)** The genomes of laboratory strains contain long stretches of reduced heterozygosity that can vary even between isolates of the supposedly same strain. In CB, observed heterozygosity is usually slightly higher than expected. The long arm of chromosome 4 has very few polymorphic sites identified in all populations, attributable to its repetitiveness. Individual data points are not indicated; lines represent loess-smoothed averages calculated from the heterozygosity of polymorphic sites. The positions of identified polymorphic sites themselves are shown for each population as a separate track at the bottom.

A clear trend emerges when one compares values for the three estimators to each other. In the wild populations, the value of *θ*_π_ is usually the smallest, followed by *θ*_w_ and then *θ*_1_ **(Figure 3A) -** a situation which is compatible with the assumption of background, or purifying, selection purging deleterious mutations from the population. In laboratory strains and in CB the order is reversed: *θ*_1_ is smallest and *θ*_π_ largest, indicating a strong demographic effect overriding the genomic signature of purifying selection.

To better understand the possible reasons for this reversed relationship we studied the inbreeding coefficient *F*_*IS*_ *= 1 - H*_*I*_*/H*_*S*_ (Wright 1951) which compares within-individual heterozygosity (*H*_*I*_) to the within-population (*H*_*S*_) heterozygosity. *F*_*IS*_ does not deviate much from zero in most wild *or* laboratory populations. The corresponding value becomes positive if multiple populations are combined. However, *F*_*IS*_ is negative in all CB substrains, which can happen only when *H*_*I*_ is larger than *H*_*S*_. This is the case when individuals are more heterozygous than expected based on the population allele frequencies, which is tantamount to a deviation from Hardy-Weinberg proportions. A possible explanation, compatible with the description of CB in the literature (Wilson, et al. 2014), is that the different CB “substrains” each result from very few, perhaps only one, pair of breeding individuals. All homozygote differences between the parents are then propagated as heterozygote differences in the children of the next few generations.

The most plausible explanation for the observed differences between TU_2014 and TU_2018, and between WIK_2014 and WIK_2018, would be a different degree of inbreeding and genetic drift. To test this, we plotted the observed and expected heterozygosity at the polymorphic sites for all strains as loess-smoothed curves across the genome (**Figure 3B, Supplementary Data S3, Supplementary Data S4**). WIK_2014 and TU_2018 were found to contain large stretches of severely reduced heterozygosity (no polymorphic sites and/or very low heterozygosity at the sites that are polymorphic), a signature of inbreeding (Kardos, et al. 2018). These patterns were less common in the TU fish from 2014 and WIK fish from 2018. In contrast, the wild fish genomes and CB do not contain such long runs of homozygosity (**Supplementary Data S3, Supplementary Data S4**). However, in the CB subgroups the observed heterozygosity in consistently higher than expected, in accordance with these being the F1 offspring of a few breeding pairs. As reported before, very few sequences can be confidently aligned to the long arm of chromosome 4 (4q), attributable to its extreme repetitiveness (Wilson, et al. 2014; Howe, et al. 2016). Therefore, very few polymorphic sites were identified on chromosome 4q from the current data. Sequencing technologies that produce longer reads than 100-300 bp would be required to tackle the variation on this part of the zebrafish genome.

### Distinct allele frequency spectra in different laboratory strains and wild populations

Frequency spectra were calculated for each population independently. This was done (1) for all SNPs, irrespective of their particular alleles **(Supplementary Data S5)**, and (2) distinguishing the two classes of A/T and G/C polymorphisms (**Figure 4**), to check for potential traces of GC-biased gene conversion (gBGC). gBGC essentially is a bias in the DNA repair machinery in favour of G or C alleles after DNA double strand breaks, for example during recombination. Although not a selection force *per se*, it can mimic the effect of selection and distort statistics that are based on the allele frequency spectrum, by increasing the frequency of G/C alleles at the expense of A/T alleles. To our knowledge, gBGC in fish has so far only been described for the three-spined stickleback (Capra and Pollard 2011)).

**Figure 4.**
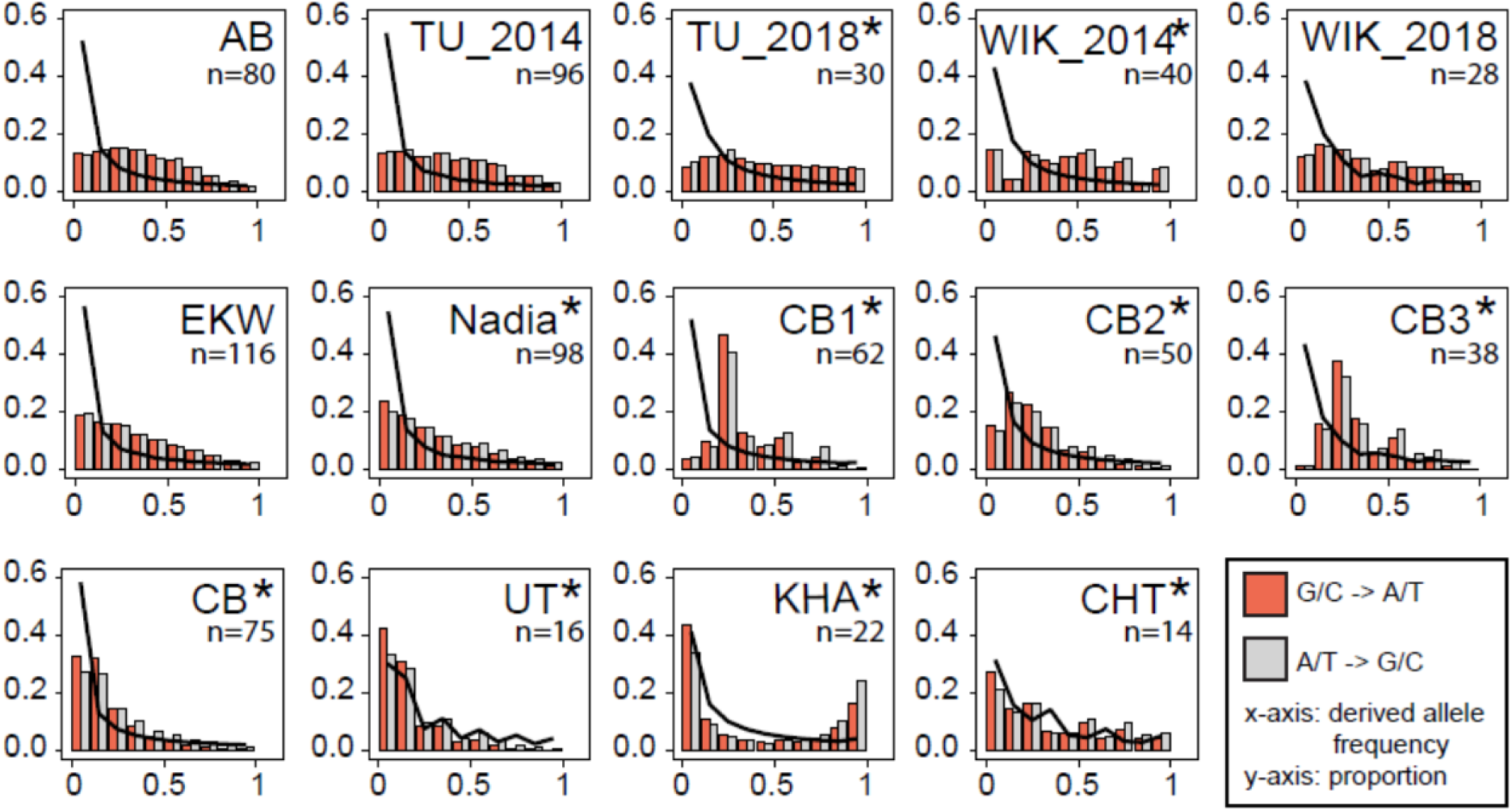
Derived allele frequency in zebrafish. **(A)** The frequency spectra in wild population closely follows the expectations (black line). In contrast, laboratory strains have a lack of low frequency alleles, with the most inbred strain (TU_2018) demonstrating a nearly flat spectrum. These spectra look similar for G/C->A/T and A/T->G/C substitution types. In the laboratory fish and the “newest” laboratory strain Nadia, biases can be seen for the substitutions that change GC-content, with A/T to G/C substitutions being more common among high frequency variants and G/C to A/T among low frequency variants. In UT and CHT the observed values are generally close to expected. The irregular shape of spectra for these two populations is caused by the uneven distribution of genomes among the bins, resulting from relatively small sample sizes. In the KHA population from Nepal, numerous alleles that are rare in other populations are present at high frequencies. Populations with significant differences between the A/T -> G/C and G/C -> A/T spectra are marked with an asterisk (*). n=number of available genomes.

In all the laboratory strains the spectra are characterized by depletion of low frequency alleles and an increase in the proportion of medium and high frequency alleles, compatible with a high amount of inbreeding and a strong decline of effective population size upon establishing laboratory strains. This effect appears to be the weakest in EKW and Nadia. Nadia is a newer strain that, at the time the sequence data were obtained, had spent only 8 generations in the laboratory (Wilson, et al. 2014) **(Figure 4)**. In the wild-derived CB there is a depletion of very low frequency alleles, which can be seen both with the three subgroups combined and separate. However, the rest of its allele frequency spectrum behaves almost as in a neutral panmictic population, i.e. proportional to the curve *1/x*. In the true wild populations the patterns are different **(Figure 4).** The frequency spectra of UT and CHT closely follow the expected power law *1/x* (times a constant). The spectrum of the KHA population from Nepal is clearly U-shaped, with a strong excess of high frequency derived alleles **(Figure 4).** Besides positive selection and local adaptation causing such a distortion of the frequency spectrum, another possible (and perhaps more likely) reason is mis-assignment of ancestral and derived status, based on the pooled consensus rule. In any case, the folded spectrum for KHA closely follows the power law and the unfolded spectrum remains U-shaped even when the reference allele is redefined based on what is the most common in the three wild zebrafish populations **(Supplementary Data S5)**.

As for gBGC, in most laboratory strains the distribution of G/C-> A/T and A/T -> G/C is similar **(Figure 4).** Chi-square test reveals significant differences between the two spectra in WIK_2014 and in TU_2018, however the spectra themselves are not consistent with gBGC. In the wild fish, as well as in the Nadia strain the p-value obtained from a chi-square test is highly significant (p < 2.2e-16) and a clear bias for G/C to A/T can be seen for low frequency alleles. For high frequency alleles, the pattern A/T to G/C is reversed, still compatible with the presence of gBGC (Glemin, et al. 2015).

In the population CB, more complicated and unusual results were observed. CB includes three subpopulations and within each one of them, there are a large number of SNPs heterozygotic in *every* individual (H_I_ = 1); up to 5% of all polymorphic sites in the (sub)population (∼3,000/60,000 for CB1 and CB3). This is in contrast to the 1-200 such sites that are seen in all the other strains (in CHT and KHA, 0.2% of all polymorphic sites). Under Hardy-Weinberg equilibrium, this is highly unlikely; for a SNP with 50% frequency, the probability of it being heterozygous in the entire sample is 1/(2^n^). Balancing selection can be also ruled out because these heterozygotic sites are evenly distributed in the genome. In addition, the number of singletons is much smaller than expected with *θ*, both in the entire CB and in subpopulations.

We propose that both phenomena are caused by the fact that CB is a F_1_ generation after a severe bottleneck, with each subpopulation deriving from one breeding pair (parents). If a SNP is homozygous for one allele in the father and homozygous for another in the mother, *all* F_1_ will be heterozygous. Under a standard equilibrium population with size N and genomic *θ* we find that the expected number of such sites (assuming that the parents are unrelated samples from the wild population) is ≈ *θ* /6, as can be calculated by the formula

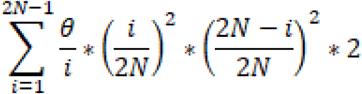

Assuming the genetic diversity of the wild parents of CB are similar to CHT, the expected number of all-heterozygote sites within the offspring of one breeding pair is ∼2,000, which is on the same scale as the actual number of 3,000. The rarity of singletons can also be explained by the fact that any SNPs must have at least an allele frequency of 25% in the parent generation (because there are only four genomes in the breeding pair) and it is unlikely that only one F_1_ genome inherited the 25% allele.

In conclusion, these results are consistent with CB being the F_1_ generation of three separate breeding pairs (three clutches of eggs). Given enough time, the descendants of CB will likely have the genetic structure of the established laboratory strains, the genomes of which contain sequence segments of high and low heterozygosity resulting from strong reduction in population size and inbreeding. (**Supplementary Data S3, Supplementary Data S4**).

## Discussion

### Differences between wild and laboratory zebrafish

We confirmed that laboratory strains have lower genetic variation than wild populations and that most (∼60%) of the variants in the dataset are only found in the wild. Because of the focus on zebrafish laboratory strains in the past, only a small fraction of the identified SNPs have been previously described. As a strain becomes more homogenous during breeding in the lab, variants become fixed or lost, which was seen from both reduced heterozygosity of the laboratory strains and the depletion of singletons and rare variants on the allele frequency spectra. The expansion of population size after the initial bottleneck causes an excess of intermediate-frequency alleles and deficit of rare ones, resulting in *θ*_π_ > *θ*_w_ > *θ*_1_. With the present dataset we could observe just how quickly this happens. For example, the AB and Nadia strains have similar proportions of polymorphic sites and nucleotide diversity. However, at the time of sample collection, Nadia had spent only eight generations in captivity and was considered a “wild-like” strain (Wilson, et al. 2014), while AB had already been established in the 1970s.

The second major difference between laboratory and wild zebrafish is in the biases in nucleotide composition that could be detected by studying the allele frequency spectra. In many species there are two opposing factors driving the nucleotide composition. The G/C -> A/T bias affects mainly alleles at low frequencies and is caused by cytosine deamination being the most common type of mutation. The A/T -> G/C bias is thought to be based on a different mechanism: during recombination, G/C rich alleles are preferred to the A/T rich alleles; the fine balance of these two mechanisms has a large impact on the genome composition. This effect can be difficult to distinguish from actual selection and has been mostly studied in mammals (Duret and Galtier 2009). However, it is also known to operate in many other taxa including land plants and bacteria (Mugal, et al. 2015). To our knowledge the GC-biased gene conversion in fish has so far only been explored in the three-spined stickleback (Capra and Pollard 2011; Roesti, et al. 2013) and never in *Cypriniformes*, the most speciose order of freshwater fishes (Nelson, et al. 2016). Here we show that the mechanism would be difficult to study in laboratory strains because GC-bias was not detectable. This can possibly be explained by the reduction of variant sites seen in the inbred laboratory strains – GC-biased gene conversion is a feature of recombination and can thus only operate on sites that have previously already become polymorphic via other mechanisms (e.g. mutation). Indeed, the bias can be observed in the Nadia strain, which was obtained more recently than the other laboratory strains of this study and has hence more polymorphic sites.

### What can we learn from the analysis of wild zebrafish?

As exemplified by the GC-biased gene conversion, it is apparent that some biological mechanisms cannot be studied in most of the laboratory strains due to their inbred nature. Among the wild zebrafish used for the current study, the Nepal KHA and Bangladesh CHT populations are less diverse than those caught or recently derived from North-East India (UT and CB). In addition, a surprisingly high number of derived alleles occurred at high frequency or were fixed in KHA. It is possible that, just like the laboratory strains, these populations have undergone a recent genetic bottleneck. Both KHA and CHT have many high frequency and fixed differences from the other fish in the dataset on both nucleotide and amino acid levels. At least some of those alleles have never been studied before, yet most likely provide the animals a selective advantage in their current habitat.

The populations in North-East India had a substantially higher proportion of polymorphic sites than any of the laboratory strains or the fish from Nepal and Bangladesh. This is likely due to a combination of mutation, gene flow, and drift. These populations are the most plausible source for those laboratory strains for which the origin is not documented, based on the results from the analysis of both genetic distances and phylogenetic trees. Indeed, major cities of the region (e.g. Kolkata) are easily accessible both by air and sea.

### Differences between laboratory strains

Differences between laboratory strains are relevant for the generality and reproducibility of zebrafish molecular studies. Fish from supposedly identical genetic backgrounds (referred to among researchers as the same strain) have been observed to provide conflicting results across different labs and different studies; supposedly minor, often undocumented details in the rearing of the fish have been pointed out as potentially impactful on the outcome of the experiments, leading to the results being difficult to reproduce (Varga, et al. 2018). Our results point to the additional explanation that genetic differences have arisen between common laboratory strains kept in different facilities. Indeed, in mice such an effect has been documented and researchers even warrant caution when interpreting results from different substrains of the widely/globally used C57BL/6 mice (Mekada, et al. 2009). We show here that fish of the same strain, but obtained from different sources and at different time points, are genetically distinct enough to be considered substrains. Although this can be seen with both TU and WIK, there are some differences in the patterns observed. TU from 2018 is the most homozygous strain in the study, whereas the TU from 2014 appears to be much more heterozygous. Conversely, WIK from 2014 appears more homozygous than WIK from 2018 and the two have fixed different amino acids in at least some proteins, which is not the case for TU.

The genetic distance between AB and the TU substrains was found to be nearly as small as the distance between the two TU substrains themselves. Considering that AB was obtained from a pet shop in the USA in 1970’s and TU from a different pet shop in Europe in the 1990’s, one possible explanation would be that they are both derived from the same undescribed pet shop strain that sold across the world over this entire period. Previous research has revealed that these two share at least one trait that distinguishes them from the other strains: they do not have an easily detectable sex determination locus at the tip of chromosome 4 (Wilson, et al. 2014). Even so, the two differ from each other in some major aspects such as structural variation (Brown, et al. 2012; Holden, et al. 2019) and haplotypes of essential immune genes, e.g. the Major Histocompatibility Complex (McConnell, et al. 2016).

While WIK and TU/AB are clearly distinct both from each other and from the wild fish, in the phylogenetic sense the populations of EKW and Nadia appear much closer to the putative source in North-East India. EKW is derived from a population maintained in Florida by commercial fish breeders; Nadia was obtained from North-East India 8 generations prior to sequencing (Wilson, et al. 2014). Compared to AB/TU/WIK, EKW and Nadia have more variants and not as flat derived allele frequency spectra, thus they can be considered more „wild-like”. However, the profile of G/C->A/T and A/T->G/C biases characteristic of wild fish is only present in Nadia and not in EKW.

In summary, this study reveals that while laboratory strains of zebrafish are an excellent model for vertebrate biology, it would be of great interest to include wild populations into more studies. They have more genetic diversity, biologic mechanisms not apparent from studying the laboratory strains (as exemplified by GC-biased gene conversion in the current study) and have distinct populations with their own preferred habitats and adaptations. Furthermore, one of the arguments for working exclusively with laboratory strains – reproducibility – does not always hold true. What is thought to be the same strain in two different laboratories may have already evolved into their own distinct substrains, as we observed for both TU and WIK in the current study.

## Materials and Methods

### Samples and datasets

The collection of wild zebrafish from India (UT), Bangladesh (CHT) and Nepal (KHA) has been described in detail by Whiteley *et al* (Whiteley, et al. 2011). The present study utilizes RAD-seq data from the same fish; a single restriction enzyme (*SbfI*) was used for library construction in order to maintain consistency with the previously published large dataset (Wilson, et al. 2014). Sequencing itself was performed on an Illumina HiSeq 2500 (2×125 bp paired end reads).

Fifteen additional fish from the TU strain and fourteen from the WIK strain were obtained in May 2018 from the Zebrafish International Resource Center in Eugene, Oregon, USA. RAD-seq libraries were produced using the protocol decribed by (Ali, et al. 2016). This protocol involves ligating biotinylated, individually indexed, adapters to RAD loci and physically separating them from the rest of the genome using streptavidin coated magnetic beads. DNA was digested with *SbfI*. Following isolation of RAD loci, Illumina sequencing adapters were added using the NEBNext Library Prep Kit for Illumina (New England Biolabs, Ipswitch, MA) using 1:10 diluted adapters and the optional size-selection step. Sequencing was preformed on an Illumina HiSeq X instrument using 2×150 bp paired end reads (Novogene Corporation Inc., Sacremento, CA).

RAD-seq data for zebrafish laboratory strains (EKW, AB, WIK, TU, Nadia), as well as for the F1 offspring of fish derived from a wild population in India (CB) were obtained from the NCBI Sequence Read Archive (Bioproject PRJNA253959). Collection of these samples, as well as generation of the sequencing data itself has been described in detail by Wilson *et al* (Wilson, et al. 2014). For downstream analyses the samples were renamed to include both the strain name and the last three characters of the SRA identifier in their labels (example: SRR1519522 became AB_522). Two fish from the original CB dataset and one from Nadia were excluded from these analyses as during test runs of the pipeline they appeared genetically different from the rest of their respective populations. For WIK, 42 fish out of 61 in the original dataset were the offspring of a single female; 41 of these were excluded from further analysis.

### Data processing

All data were generated by sequencing zebrafish genomic DNA digested with the enzyme *SbfI*. Wilson *et al* sequences (Wilson, et al. 2014) originate from single-end sequencing and have a length of 95bp. For compatibility, all raw data from the newly sequenced populations was first trimmed to the same length (this was done with GNU cut), then de-multiplexed and cleaned using *process_radtags* from Stacks v2.4 (Catchen, et al. 2011; Rochette and Catchen 2017). Sequences were mapped to the zebrafish reference genome (GRCz11; downloaded from Ensembl) using *bwa mem* (version 0.7.17-r1188) (Li and Durbin 2009) with the default settings. *Samtools* (version 1.9) was used to filter out unmapped reads and non-primary alignments. Variant calling was performed with *Stacks v2.4* (Catchen, et al. 2011; Rochette and Catchen 2017). In order to deal with possible mutations of the cutting site and subsequent allele dropout, data were considered for analysis only if available from at least 70% of populations, and in at least 80% of the individuals within each population. The log likelihood threshold for Stacks to retain RAD loci was -10.

### Analysis of population structure

Admixture analysis of the population structure was performed with the R package *LEA* (Frichot and Francois 2015), using a single SNP per RAD locus. The optimal number of populations was chosen based on minimal entropy analysis performed by the software. Concatenated sequences of variant positions in the data (one sequence per population; encoded according to the IUPAC nomenclature) were used as input for Maximum Likelihood tree construction with *RAxML v8.2.12* (Stamatakis 2014), using the GTRCAT model and 1,000 bootstrap replicates. Estimates for genetic distances were obtained from the *populations* module of *Stacks*.

### Data analysis

Estimates of heterozygosity, nucleotide diversity and population differentiation were obtained from the *populations* module of *Stacks* (Catchen, et al. 2011). Variant sites were annotated with the *Ensembl Variant Effect Predictor (VEP)* (McLaren, et al. 2016).

We determined three estimates of the scaled mutation rate *θ=4Nμ*, Tajima’s estimator *θ*_*π*_ (Tajima 1989), Watterson’s estimator *θ* _*w*_ (Watterson 1975) and *θ*_*1*_, an estimator of *θ* based only on singleton mutations (Fu and Li 1993). The first one is the average number of nucleotide differences in pairwise comparisons, and the second is obtained by dividing the total number of polymorphic sites within a sample by *h*_*2n-1*_, the harmonic number of the (diploid) sample size minus one. All estimators are rescaled to values per bp.

Allele frequencies were extracted from the *stacks*-generated .vcf (Variant Call Format) file with *VCFtools* (version 0.1.13) (Danecek, et al. 2011). *Stacks* assumes data to be biallelic and by default considers the less frequent allele in the dataset to be derived. These data were used to plot allele frequency spectra and to extract differentially fixed alleles. Expected allele frequencies were calculated assuming a power law distribution of 1/x * constant. Both the observed and expected values were summed to bins of 10% frequency before plotting. In parallel, the same was performed when redefining the ‘ancestral’ and ‘derived’ alleles based on what is the most common in a reduced dataset of seven fish each from the three major wild lineages – India, Nepal and Bangladesh **(Supplementary Data S5).**

### Data visualization and statistics

R core tools and the R packages *gplots* and *ggplot2* were used to generate the initial plots (Warnes, et al. 2016; Wickham 2016; R Core Team 2018). These were then imported to Adobe Illustrator CS6 (version 16.03) for final editing. Chi-square tests for the allele frequency distributions of each population were performed with R.

## Supporting information

Supplemental Data S1 - heatmaps

Supplemental Data S2 - fixed nonsynonymous differences

Supplemental Data S3 - heterozygosity chr1-12

Supplemental Data S4 - heterozygosity chr13-25

Supplemental Data S5 - frequency spectra

## Data Availability

All generated sequence data is available for download from the NCBI Sequence Read Archive (SRA); BioProject PRJNA555030. The scripts used in this study are available from GitHub at https://github.com/jsuurvali/rad_analysis. Files containing the identified variant sites and their functional annotations are available from the Dryad Digital Repository <placeholder_for_ID>.

## Author Contributions

JS supervised sequencing, analysed the data and wrote the manuscript. AW contributed sample material and RAD-sequencing of laboratory zebrafish. KG performed RAD-sequencing of the wild zebrafish. TW and YZ contributed to data analysis. ML and TW conceived the study and revised the manuscript.

## Acknowledgements

This study was supported by the German Research Council (DFG) grant SPP1819 to TW and ML. AW was supported by the National Science Foundation award DEB-1652278 and by a subaward to the National Science Foundation award IOS-1257562. Emilia Martins, Anuradha Bhat, Rick Mayden and Jiwan Shrestha helped obtain the wild zebrafish samples reported in this paper. Seth Smith helped with the design and implementation of the RAD-Seq protocols used to obtain sequence data for TU and WIK. Kamel Jabbari provided helpful feedback and advice for the analysis of GC-biased gene conversion.

