## Supplemental Data S1 - heatmaps for "The laboratory domestication of zebrafish: from diverse populations to inbred substrains"

Supplementary data S1 for:

### The laboratory domestication of zebrafish: from diverse populations to inbred substrains

Jaanus Suurväli, Andrew R. Whiteley, Yichen Zheng, Karim Gharbi, Maria Leptin, Thomas Wiehe

#### Heatmaps for measures of genetic differentiation between different wild and laboratory zebrafish populations

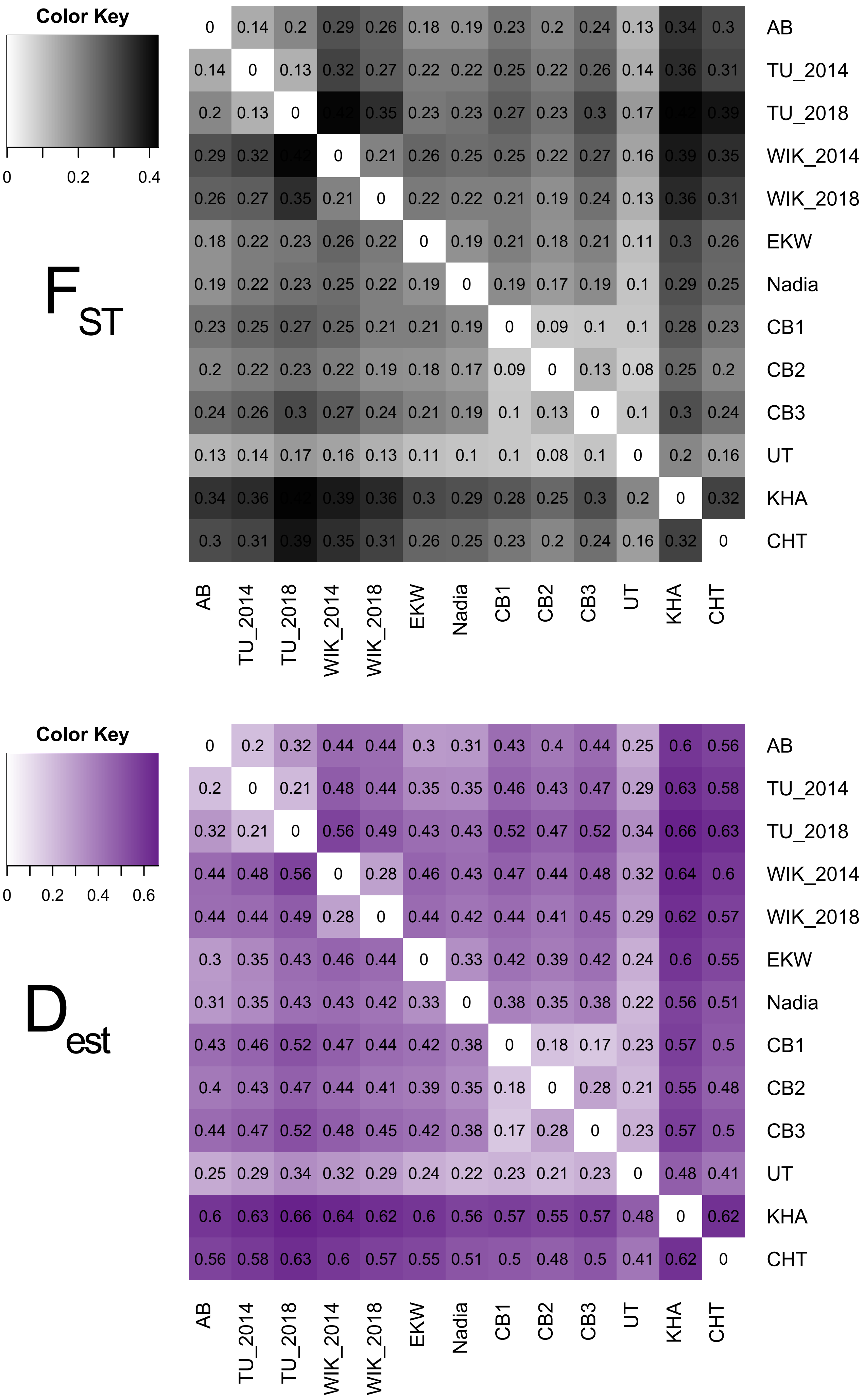
Color Key
0
0.2
0.4
FST
AB 0 0.14 0.2 0.29 0.26 0.18 0.19 0.23 0.2 0.24 0.13 0.34 0.3
TU_2014 0.14 0 0.13 0.32 0.27 0.22 0.22 0.25 0.22 0.26 0.14 0.36 0.31
TU_2018 0.2 0.13 0 0.42 0.35 0.23 0.23 0.27 0.23 0.3 0.17 0.42 0.39
WIK_2014 0.29 0.32 0.42 0 0.21 0.26 0.25 0.25 0.22 0.27 0.16 0.39 0.35
WIK_2018 0.26 0.27 0.35 0.21 0 0.22 0.22 0.21 0.19 0.24 0.13 0.36 0.31
EKW 0.18 0.22 0.23 0.26 0.22 0 0.19 0.21 0.18 0.21 0.11 0.3 0.26
Nadia 0.19 0.22 0.23 0.25 0.22 0.19 0 0.19 0.17 0.19 0.1 0.29 0.25
CB1 0.23 0.25 0.27 0.25 0.21 0.21 0.19 0 0.09 0.1 0.1 0.28 0.23
CB2 0.2 0.22 0.23 0.22 0.19 0.18 0.17 0.09 0 0.13 0.08 0.25 0.2
CB3 0.24 0.26 0.3 0.27 0.24 0.21 0.19 0.1 0.13 0 0.1 0.3 0.24
UT 0.13 0.14 0.17 0.16 0.13 0.11 0.1 0.1 0.08 0.1 0 0.2 0.16
KHA 0.34 0.36 0.42 0.39 0.36 0.3 0.29 0.28 0.25 0.3 0.2 0 0.32
CHT 0.3 0.31 0.39 0.35 0.31 0.26 0.25 0.23 0.2 0.24 0.16 0.32 0
AB TU_2014 TU_2018 WIK_2014 WIK_2018 EKW Nadia CB1 CB2 CB3 UT KHA CHT
Color Key
0
0.2
0.4
0.6
Dest
AB 0 0.2 0.32 0.44 0.44 0.3 0.31 0.43 0.4 0.44 0.25 0.6 0.56
TU_2014 0.2 0 0.21 0.48 0.44 0.35 0.35 0.46 0.43 0.47 0.29 0.63 0.58
TU_2018 0.32 0.21 0 0.56 0.49 0.43 0.43 0.52 0.47 0.52 0.34 0.66 0.63
WIK_2014 0.44 0.48 0.56 0 0.28 0.46 0.43 0.47 0.44 0.48 0.32 0.64 0.6
WIK_2018 0.44 0.44 0.49 0.28 0 0.44 0.42 0.44 0.41 0.45 0.29 0.62 0.57
EKW 0.3 0.35 0.43 0.46 0.44 0 0.33 0.42 0.39 0.42 0.24 0.6 0.55
Nadia 0.31 0.35 0.43 0.43 0.42 0.33 0 0.38 0.35 0.38 0.22 0.56 0.51
CB1 0.43 0.46 0.52 0.47 0.44 0.42 0.38 0 0.18 0.17 0.23 0.57 0.5
CB2 0.4 0.43 0.47 0.44 0.41 0.39 0.35 0.18 0 0.28 0.21 0.55 0.48
CB3 0.44 0.47 0.52 0.48 0.45 0.42 0.38 0.17 0.28 0 0.23 0.57 0.5
UT 0.25 0.29 0.34 0.32 0.29 0.24 0.22 0.23 0.21 0.23 0 0.48 0.41
KHA 0.6 0.63 0.66 0.64 0.62 0.6 0.56 0.57 0.55 0.57 0.48 0 0.62
CHT 0.56 0.58 0.63 0.6 0.57 0.55 0.51 0.5 0.48 0.5 0.41 0.62 0
AB TU_2014 TU_2018 WIK_2014 WIK_2018 EKW Nadia CB1 CB2 CB3 UT KHA CHT

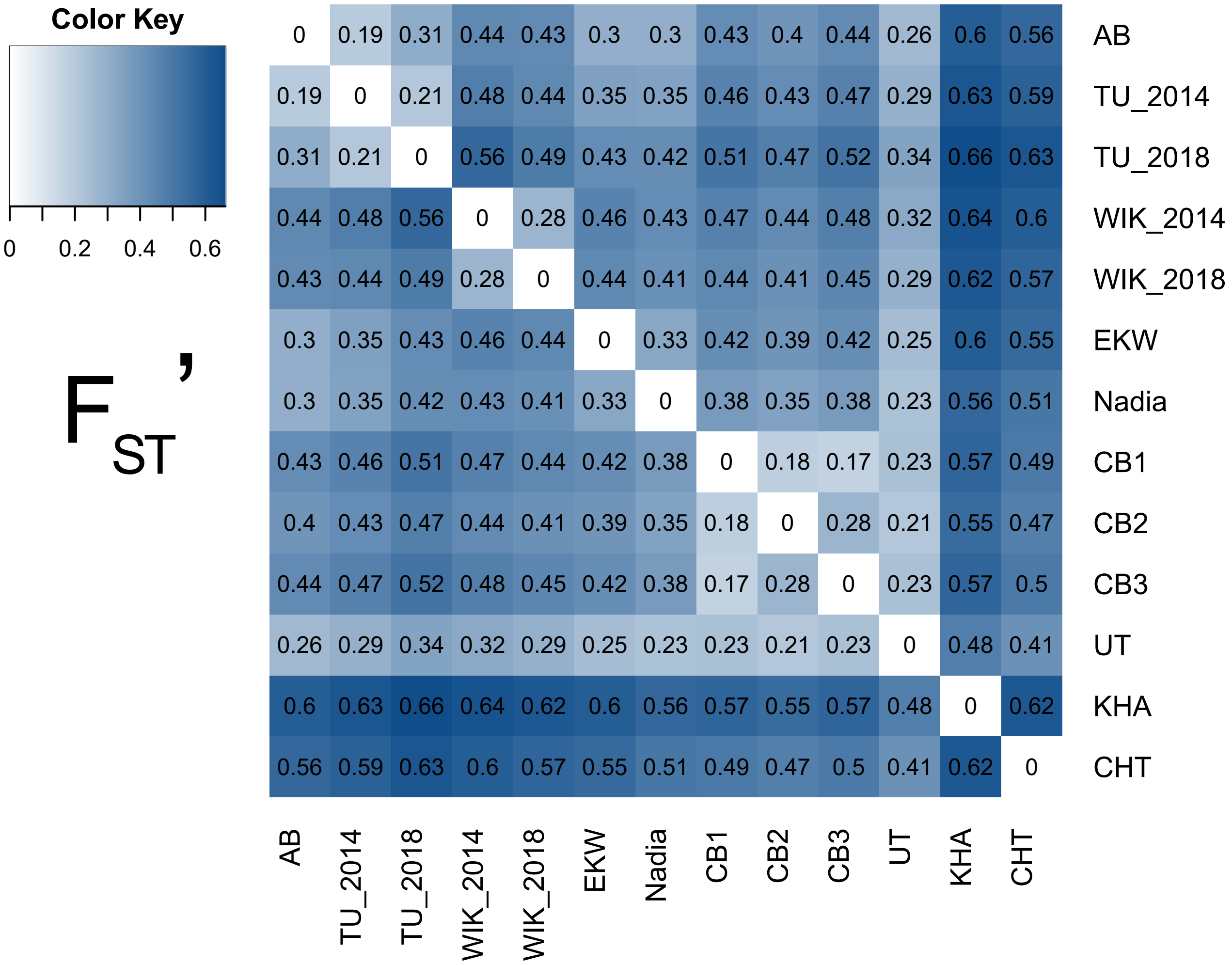
Color Key
0 0.2 0.4 0.6
$F_{ST}'$
0 0.19 0.31 0.44 0.43 0.3 0.3 0.43 0.4 0.44 0.26 0.6 0.56 AB
0.19 0 0.21 0.48 0.44 0.35 0.35 0.46 0.43 0.47 0.29 0.63 0.59 TU_2014
0.31 0.21 0 0.56 0.49 0.43 0.42 0.51 0.47 0.52 0.34 0.66 0.63 TU_2018
0.44 0.48 0.56 0 0.28 0.46 0.43 0.47 0.44 0.48 0.32 0.64 0.6 WIK_2014
0.43 0.44 0.49 0.28 0 0.44 0.41 0.44 0.41 0.45 0.29 0.62 0.57 WIK_2018
0.3 0.35 0.43 0.46 0.44 0 0.33 0.42 0.39 0.42 0.25 0.6 0.55 EKW
0.3 0.35 0.42 0.43 0.41 0.33 0 0.38 0.35 0.38 0.23 0.56 0.51 Nadia
0.43 0.46 0.51 0.47 0.44 0.42 0.38 0 0.18 0.17 0.23 0.57 0.49 CB1
0.4 0.43 0.47 0.44 0.41 0.39 0.35 0.18 0 0.28 0.21 0.55 0.47 CB2
0.44 0.47 0.52 0.48 0.45 0.42 0.38 0.17 0.28 0 0.23 0.57 0.5 CB3
0.26 0.29 0.34 0.32 0.29 0.25 0.23 0.23 0.21 0.23 0 0.48 0.41 UT
0.6 0.63 0.66 0.64 0.62 0.6 0.56 0.57 0.55 0.57 0.48 0 0.62 KHA
0.56 0.59 0.63 0.6 0.57 0.55 0.51 0.49 0.47 0.5 0.41 0.62 0 CHT
AB TU_2014 TU_2018 WIK_2014 WIK_2018 EKW Nadia CB1 CB2 CB3 UT KHA CHT

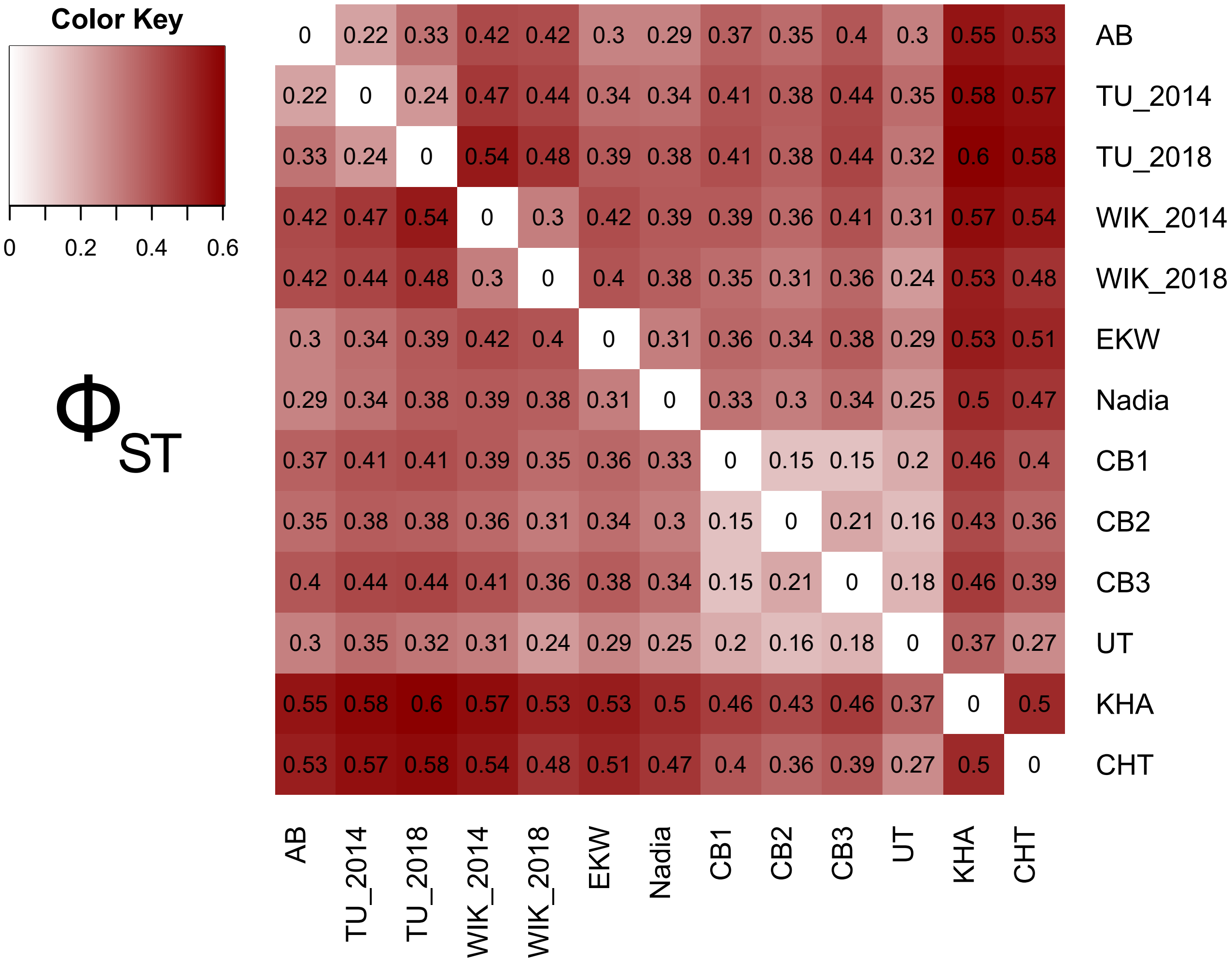
Color Key
0 0.2 0.4 0.6
$\Phi_{ST}$
0 0.22 0.33 0.42 0.42 0.3 0.29 0.37 0.35 0.4 0.3 0.55 0.53 AB
0.22 0 0.24 0.47 0.44 0.34 0.34 0.41 0.38 0.44 0.35 0.58 0.57 TU_2014
0.33 0.24 0 0.54 0.48 0.39 0.38 0.41 0.38 0.44 0.32 0.6 0.58 TU_2018
0.42 0.47 0.54 0 0.3 0.42 0.39 0.39 0.36 0.41 0.31 0.57 0.54 WIK_2014
0.42 0.44 0.48 0.3 0 0.4 0.38 0.35 0.31 0.36 0.24 0.53 0.48 WIK_2018
0.3 0.34 0.39 0.42 0.4 0 0.31 0.36 0.34 0.38 0.29 0.53 0.51 EKW
0.29 0.34 0.38 0.39 0.38 0.31 0 0.33 0.3 0.34 0.25 0.5 0.47 Nadia
0.37 0.41 0.41 0.39 0.35 0.36 0.33 0 0.15 0.15 0.2 0.46 0.4 CB1
0.35 0.38 0.38 0.36 0.31 0.34 0.3 0.15 0 0.21 0.16 0.43 0.36 CB2
0.4 0.44 0.44 0.41 0.36 0.38 0.34 0.15 0.21 0 0.18 0.46 0.39 CB3
0.3 0.35 0.32 0.31 0.24 0.29 0.25 0.2 0.16 0.18 0 0.37 0.27 UT
0.55 0.58 0.6 0.57 0.53 0.53 0.5 0.46 0.43 0.46 0.37 0 0.5 KHA
0.53 0.57 0.58 0.54 0.48 0.51 0.47 0.4 0.36 0.39 0.27 0.5 0 CHT
AB TU_2014 TU_2018 WIK_2014 WIK_2018 EKW Nadia CB1 CB2 CB3 UT KHA CHT
