## Supplemental Data S3 - heterozygosity chr1-12 for "The laboratory domestication of zebrafish: from diverse populations to inbred substrains"

Supplementary data S3/S4 for:

### The laboratory domestication of zebrafish: from diverse populations to inbred substrains

Jaanus Suurväli, Andrew R. Whiteley, Yichen Zheng, Karim Gharbi, Maria Leptin, Thomas Wiehe

Heterozygosity in wild and laboratory zebrafish.

Data was smoothed with Loess regression (span=0.06) prior to plotting it across chromosomes.

Supplementary Data S3: chromosomes 1-12

Supplementary Data S4: chromosomes 13-25

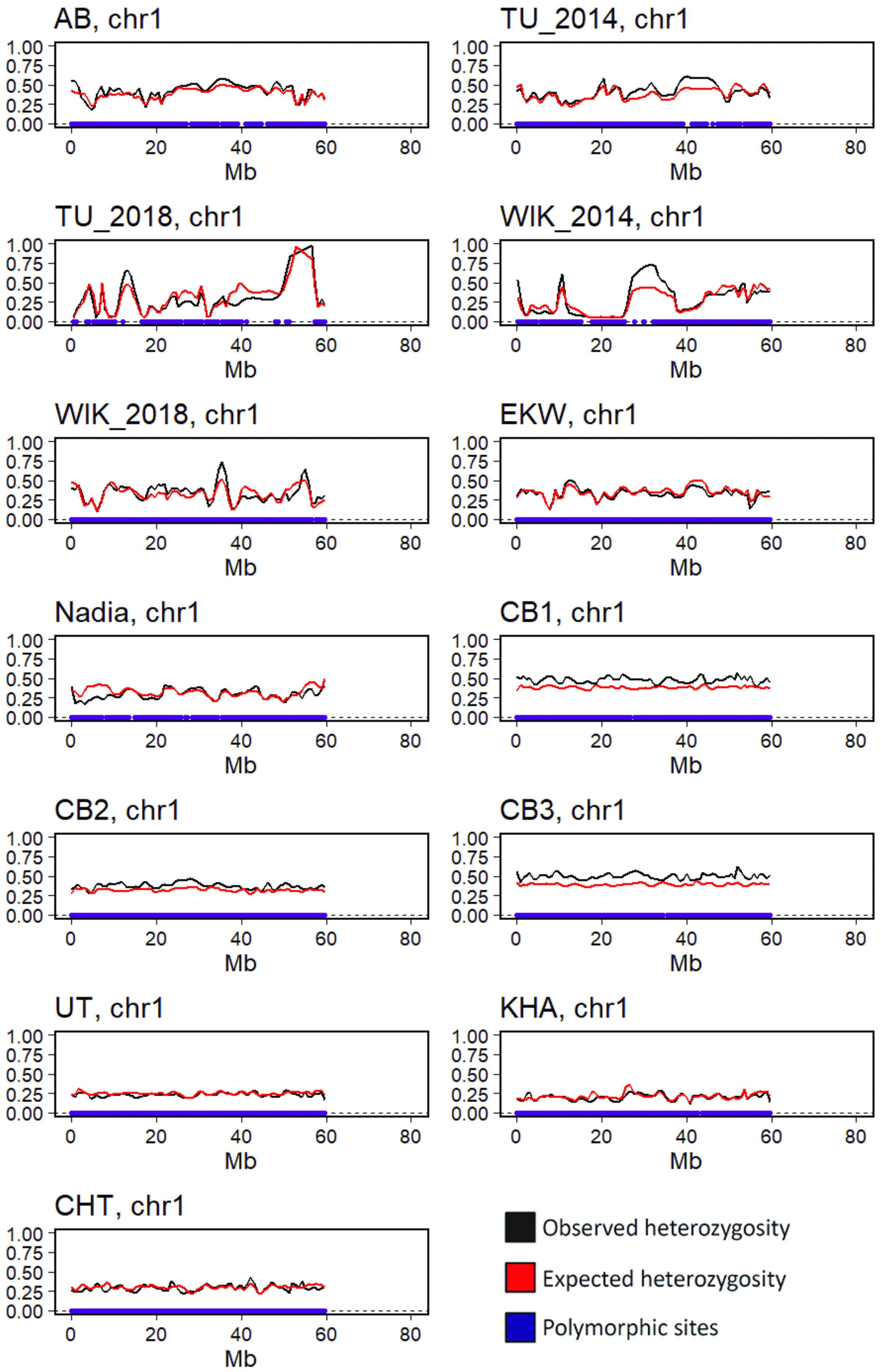
AB, chr1
TU_2014, chr1
TU_2018, chr1
WIK_2014, chr1
WIK_2018, chr1
EKW, chr1
Nadia, chr1
CB1, chr1
CB2, chr1
CB3, chr1
UT, chr1
KHA, chr1
CHT, chr1
1.00
0.75
0.50
0.25
0.00
0
20
40
60
80
Mb
Observed heterozygosity
Expected heterozygosity
Polymorphic sites

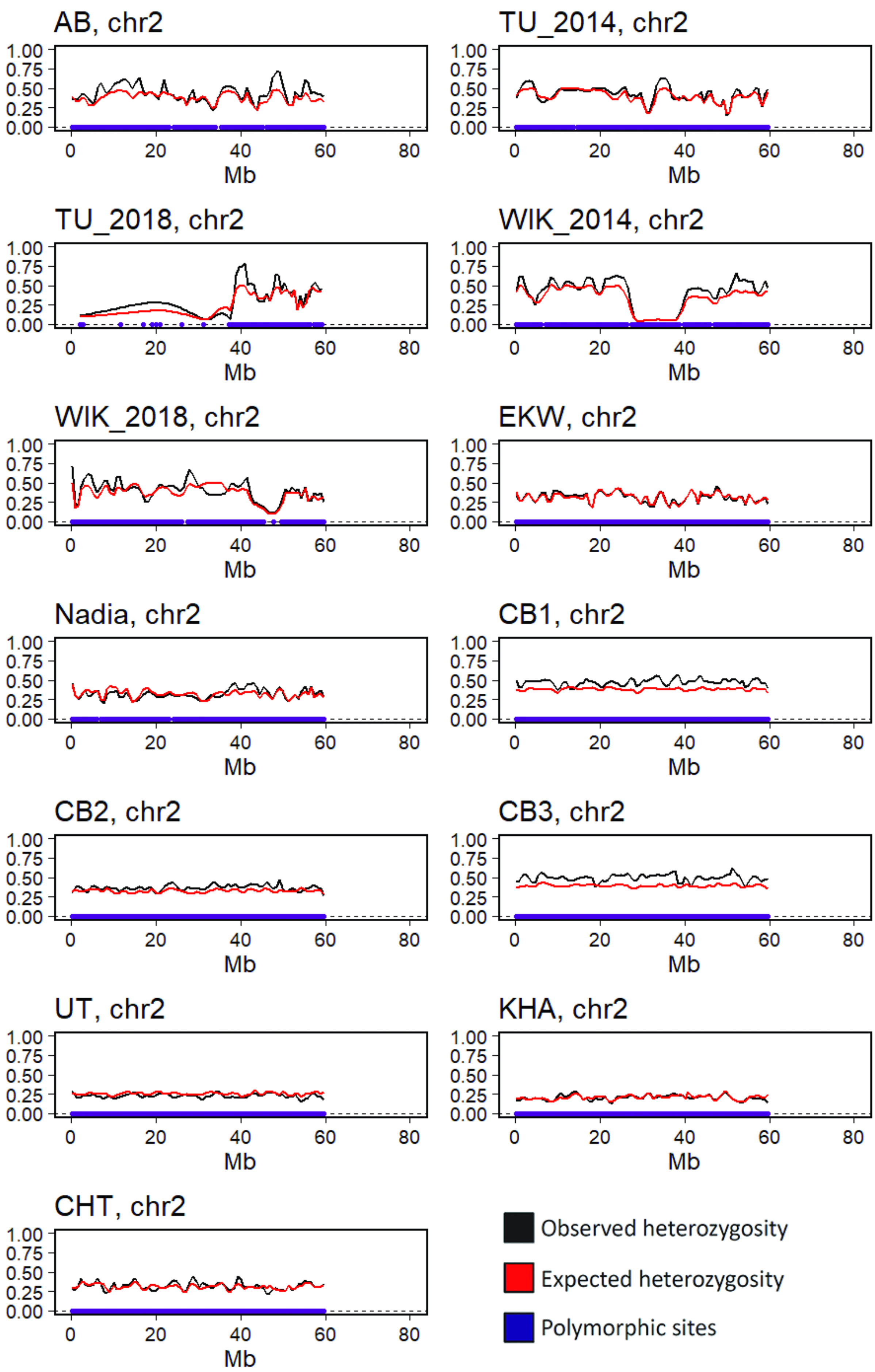
AB, chr2
TU_2014, chr2
TU_2018, chr2
WIK_2014, chr2
WIK_2018, chr2
EKW, chr2
Nadia, chr2
CB1, chr2
CB2, chr2
CB3, chr2
UT, chr2
KHA, chr2
CHT, chr2
1.00
0.75
0.50
0.25
0.00
0
20
40
60
80
Mb
Observed heterozygosity
Expected heterozygosity
Polymorphic sites

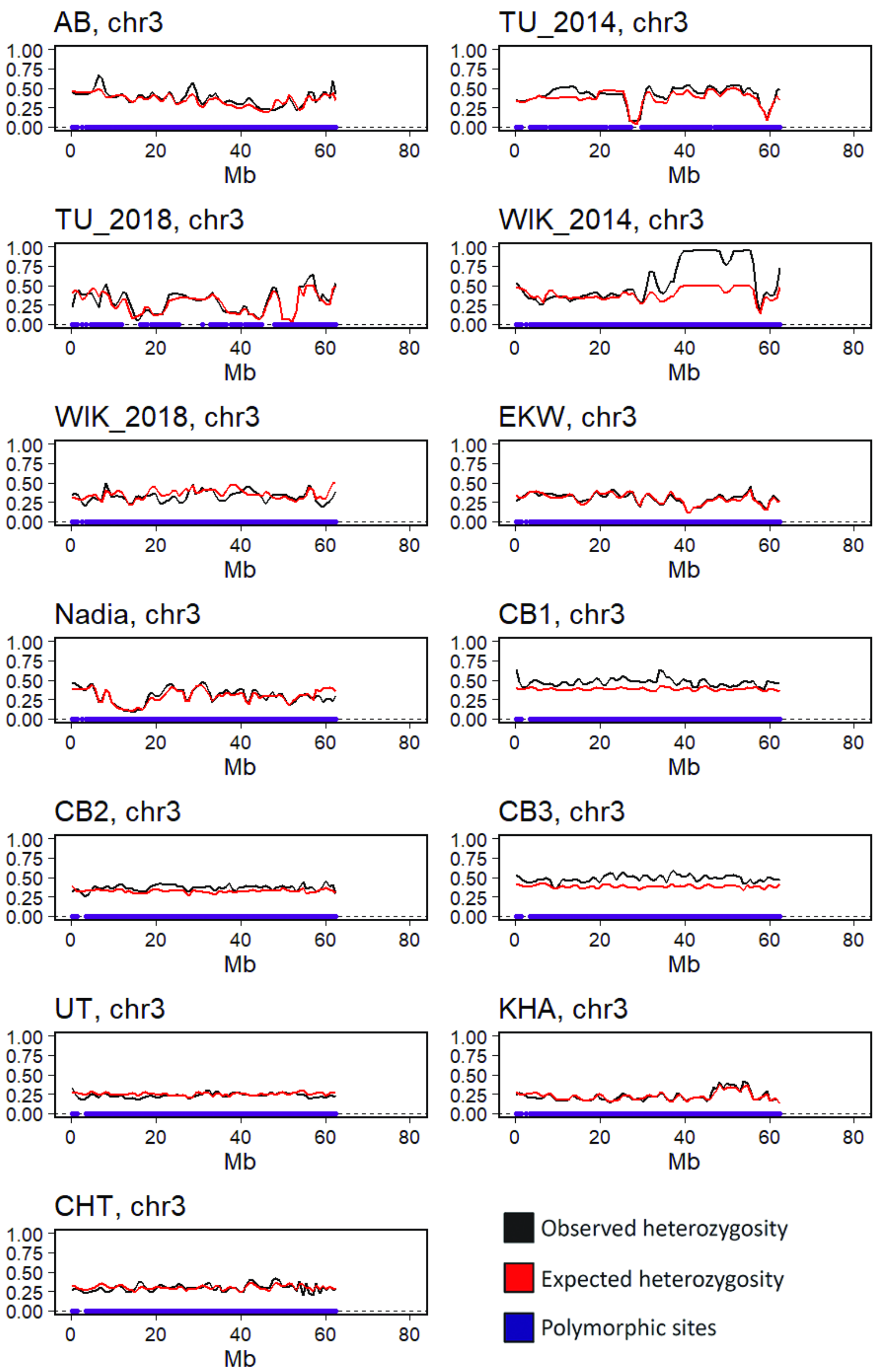
AB, chr3
TU_2014, chr3
TU_2018, chr3
WIK_2014, chr3
WIK_2018, chr3
EKW, chr3
Nadia, chr3
CB1, chr3
CB2, chr3
CB3, chr3
UT, chr3
KHA, chr3
CHT, chr3
1.00
0.75
0.50
0.25
0.00
0
20
40
60
80
Mb
Observed heterozygosity
Expected heterozygosity
Polymorphic sites

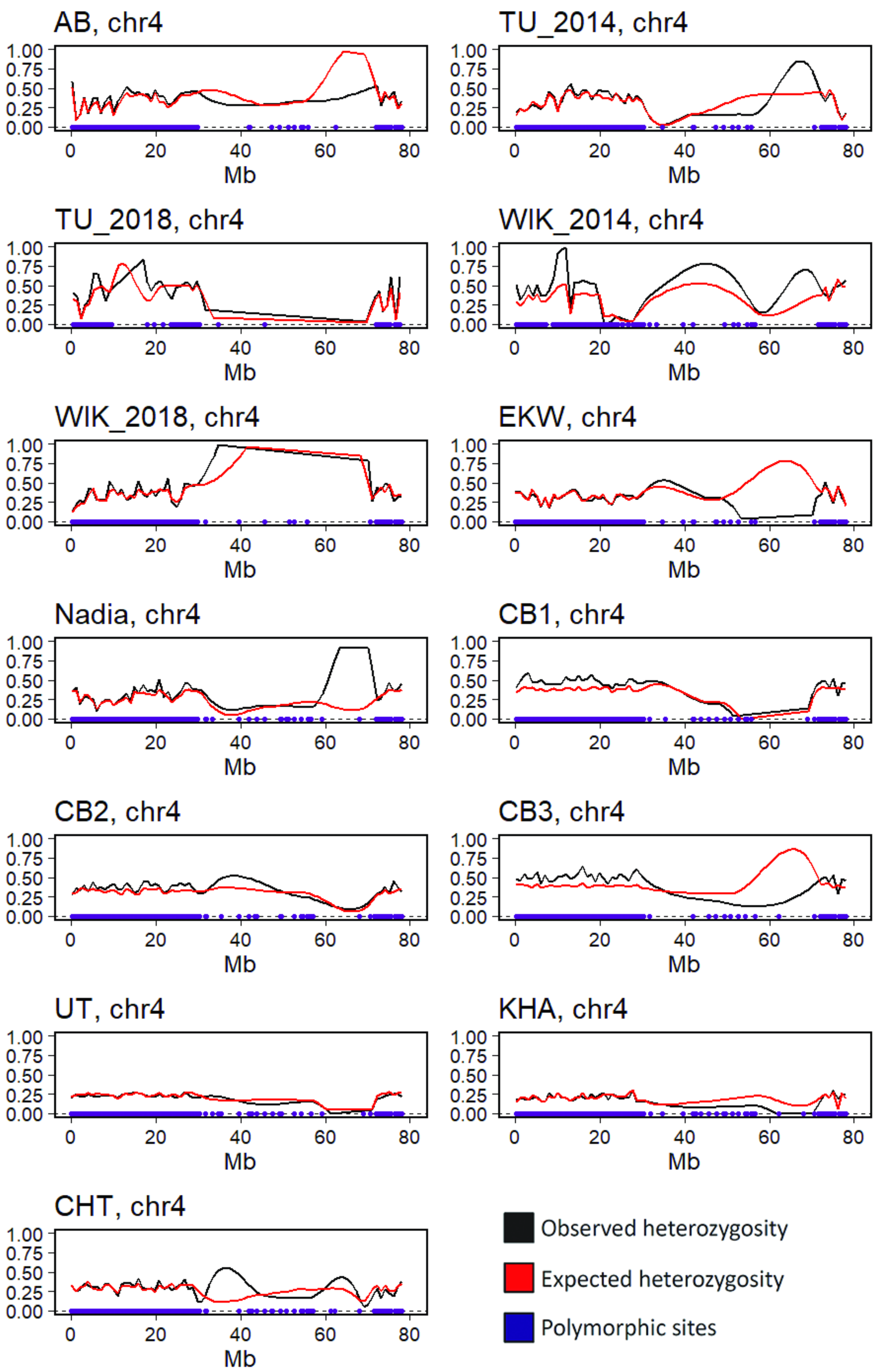
AB, chr4
TU_2014, chr4
TU_2018, chr4
WIK_2014, chr4
WIK_2018, chr4
EKW, chr4
Nadia, chr4
CB1, chr4
CB2, chr4
CB3, chr4
UT, chr4
KHA, chr4
CHT, chr4
1.00
0.75
0.50
0.25
0.00
0
20
40
60
80
Mb
Observed heterozygosity
Expected heterozygosity
Polymorphic sites

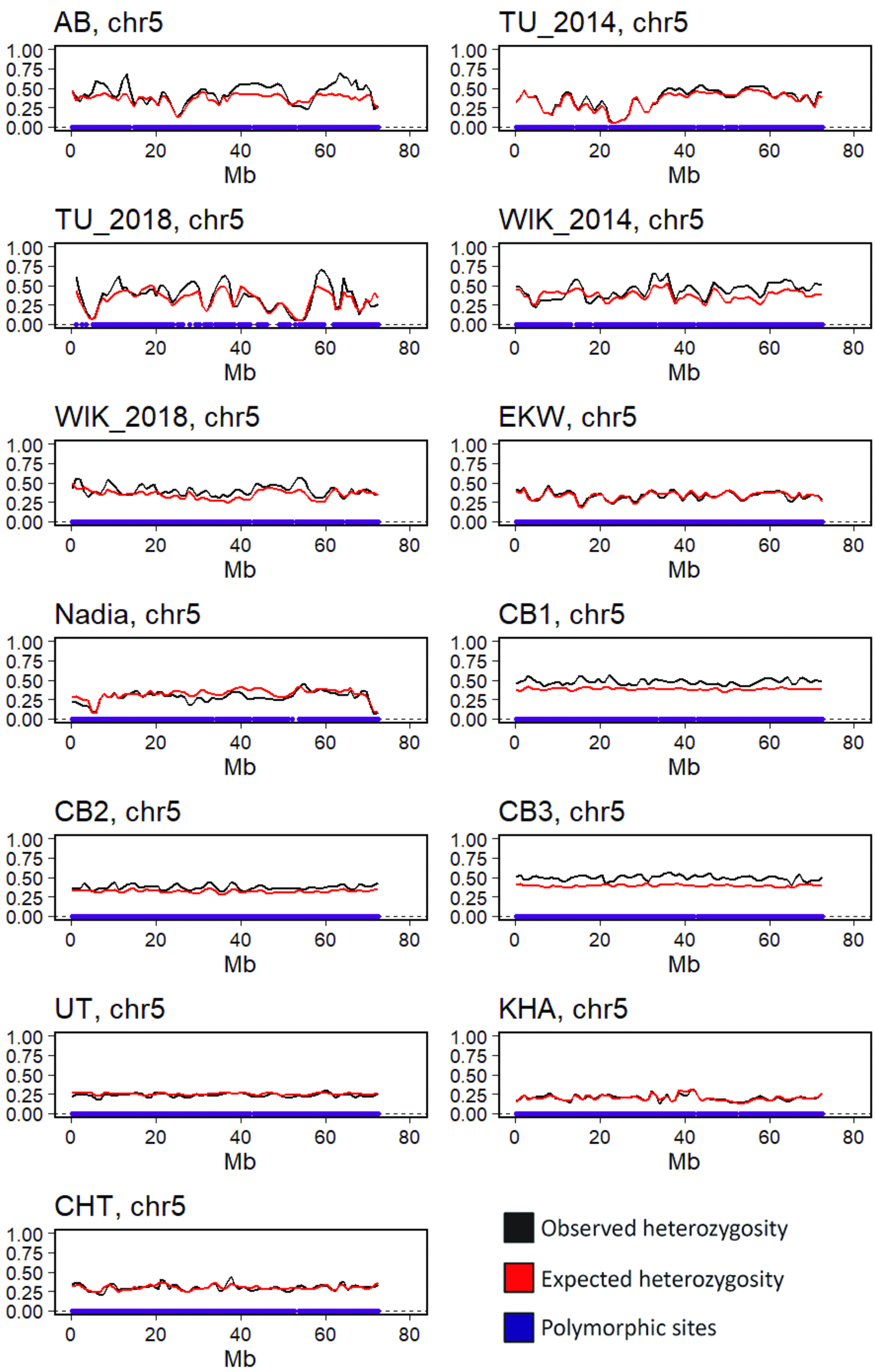
AB, chr5
TU_2014, chr5
TU_2018, chr5
WIK_2014, chr5
WIK_2018, chr5
EKW, chr5
Nadia, chr5
CB1, chr5
CB2, chr5
CB3, chr5
UT, chr5
KHA, chr5
CHT, chr5
1.00
0.75
0.50
0.25
0.00
0
20
40
60
80
Mb
Observed heterozygosity
Expected heterozygosity
Polymorphic sites

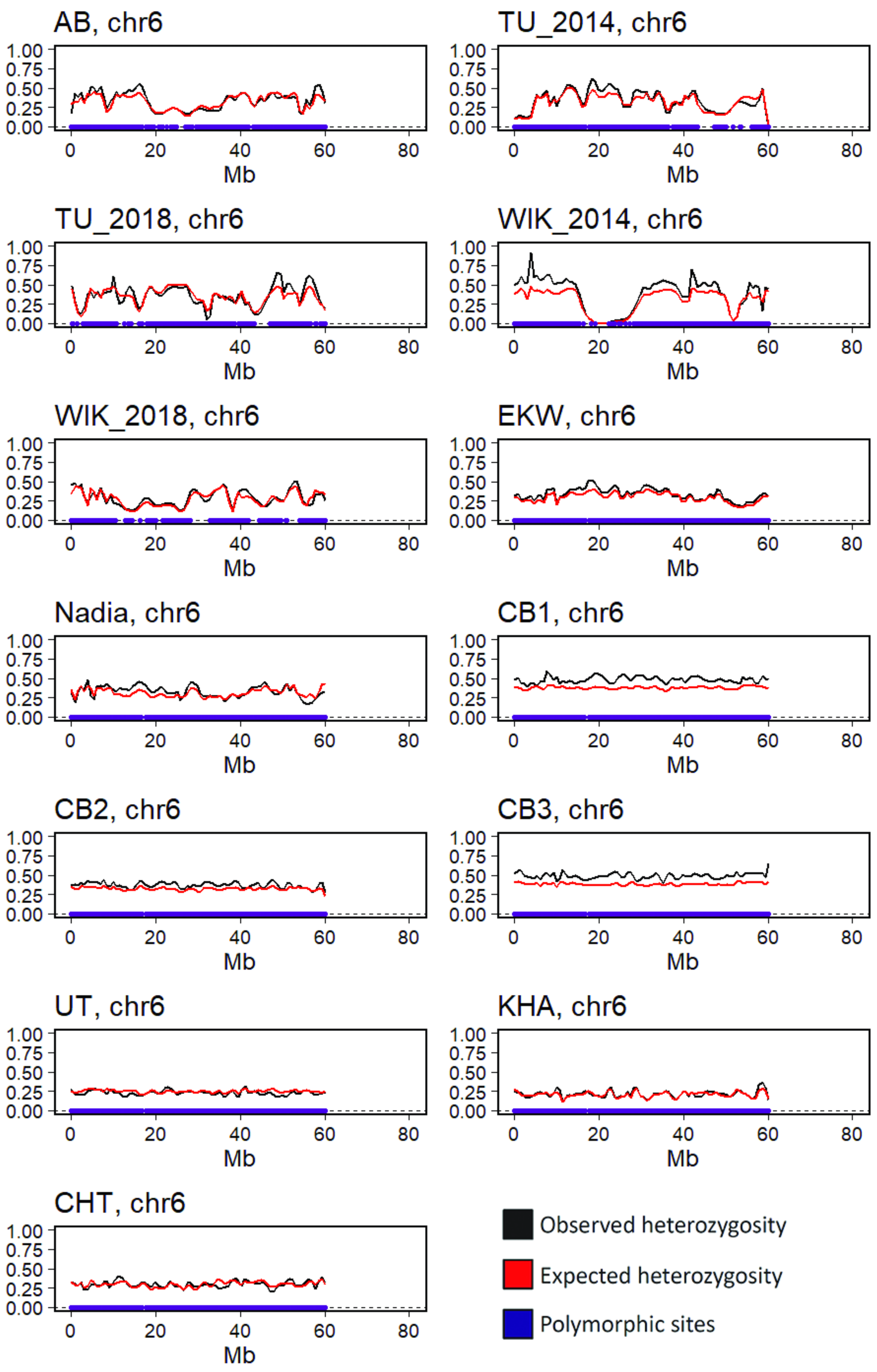

AB, chr6
TU_2014, chr6
TU_2018, chr6
WIK_2014, chr6
WIK_2018, chr6
EKW, chr6
Nadia, chr6
CB1, chr6
CB2, chr6
CB3, chr6
UT, chr6
KHA, chr6
CHT, chr6
1.00
0.75
0.50
0.25
0.00
0
20
40
60
80
Mb
Observed heterozygosity
Expected heterozygosity
Polymorphic sites

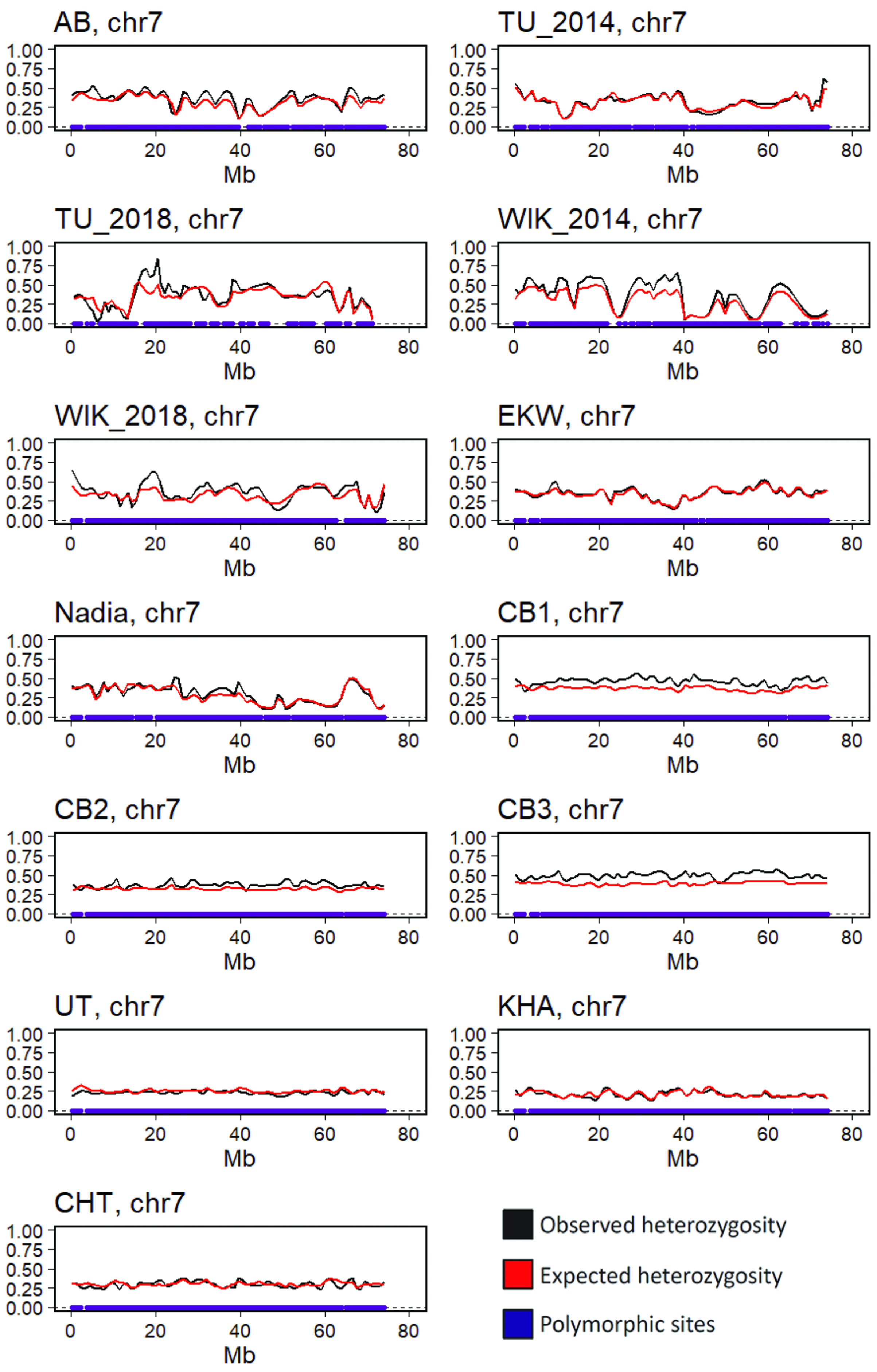
AB, chr7
TU_2014, chr7
TU_2018, chr7
WIK_2014, chr7
WIK_2018, chr7
EKW, chr7
Nadia, chr7
CB1, chr7
CB2, chr7
CB3, chr7
UT, chr7
KHA, chr7
CHT, chr7
1.00
0.75
0.50
0.25
0.00
0
20
40
60
80
Mb
Observed heterozygosity
Expected heterozygosity
Polymorphic sites

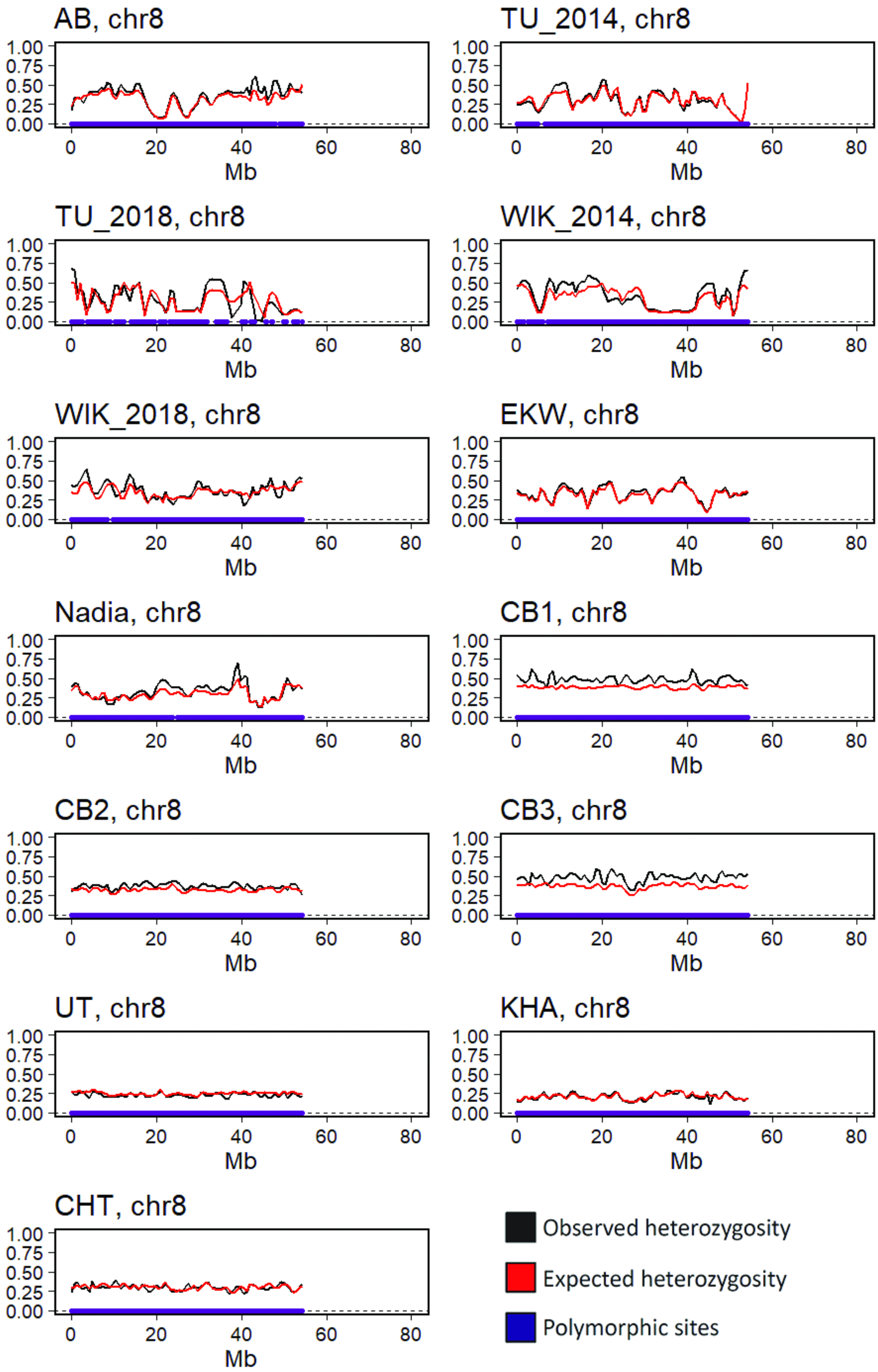

AB, chr8
TU_2014, chr8
TU_2018, chr8
WIK_2014, chr8
WIK_2018, chr8
EKW, chr8
Nadia, chr8
CB1, chr8
CB2, chr8
CB3, chr8
UT, chr8
KHA, chr8
CHT, chr8
1.00
0.75
0.50
0.25
0.00
0
20
40
60
80
Mb
Observed heterozygosity
Expected heterozygosity
Polymorphic sites

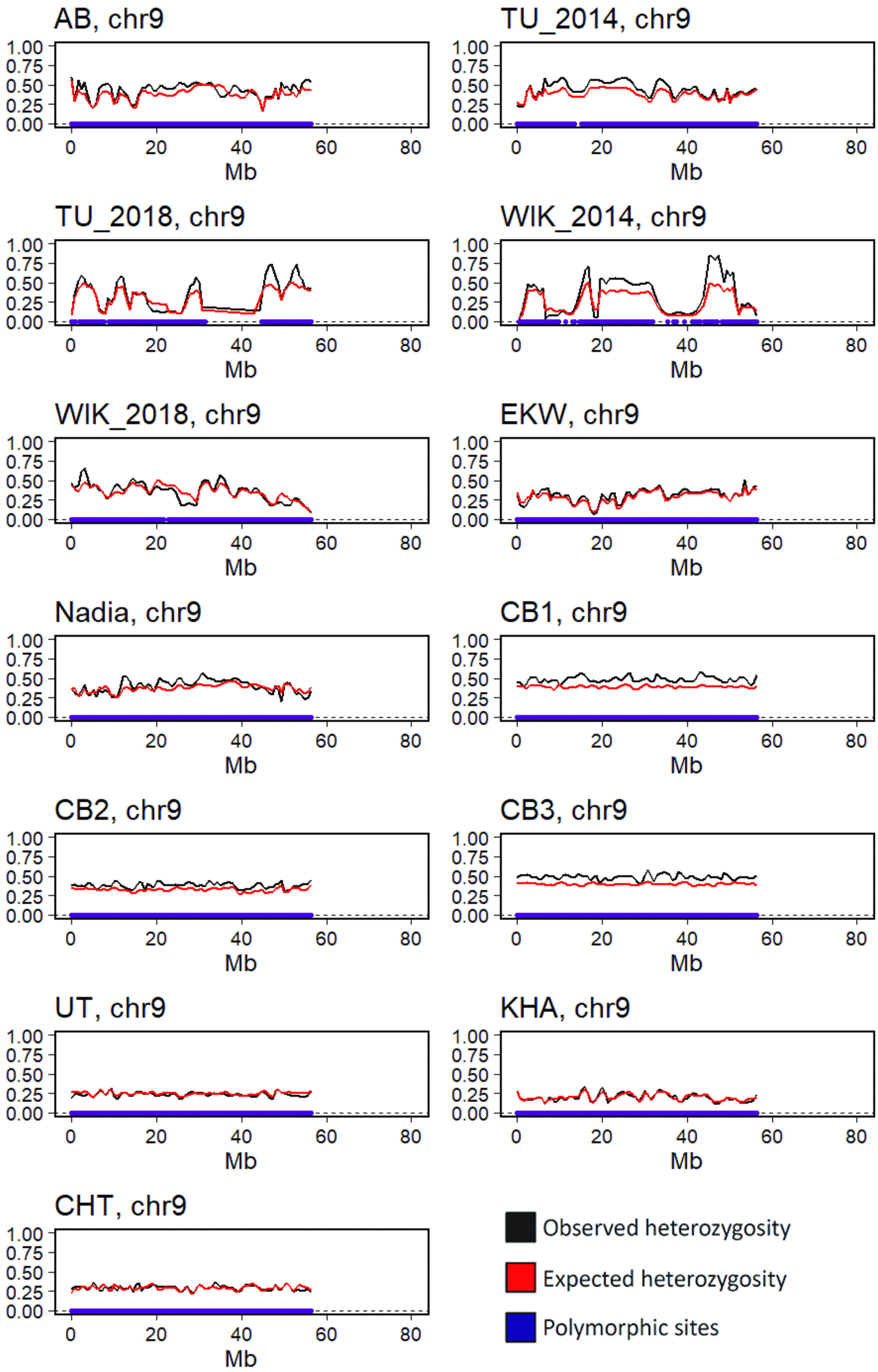
AB, chr9
TU_2014, chr9
TU_2018, chr9
WIK_2014, chr9
WIK_2018, chr9
EKW, chr9
Nadia, chr9
CB1, chr9
CB2, chr9
CB3, chr9
UT, chr9
KHA, chr9
CHT, chr9
1.00
0.75
0.50
0.25
0.00
0
20
40
60
80
Mb
Observed heterozygosity
Expected heterozygosity
Polymorphic sites

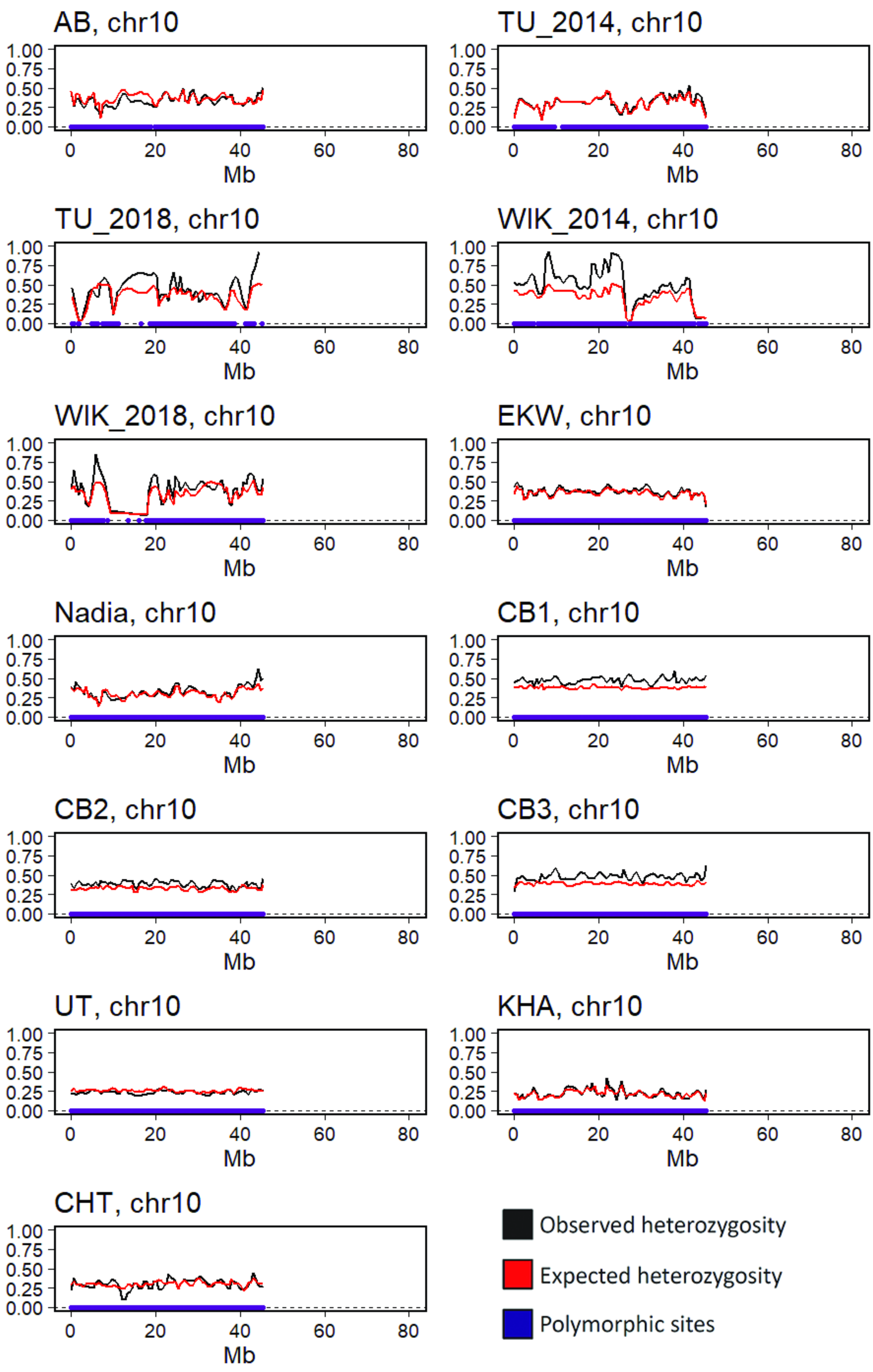
AB, chr10
TU_2014, chr10
TU_2018, chr10
WIK_2014, chr10
WIK_2018, chr10
EKW, chr10
Nadia, chr10
CB1, chr10
CB2, chr10
CB3, chr10
UT, chr10
KHA, chr10
CHT, chr10
1.00
0.75
0.50
0.25
0.00
0
20
40
60
80
Mb
Observed heterozygosity
Expected heterozygosity
Polymorphic sites

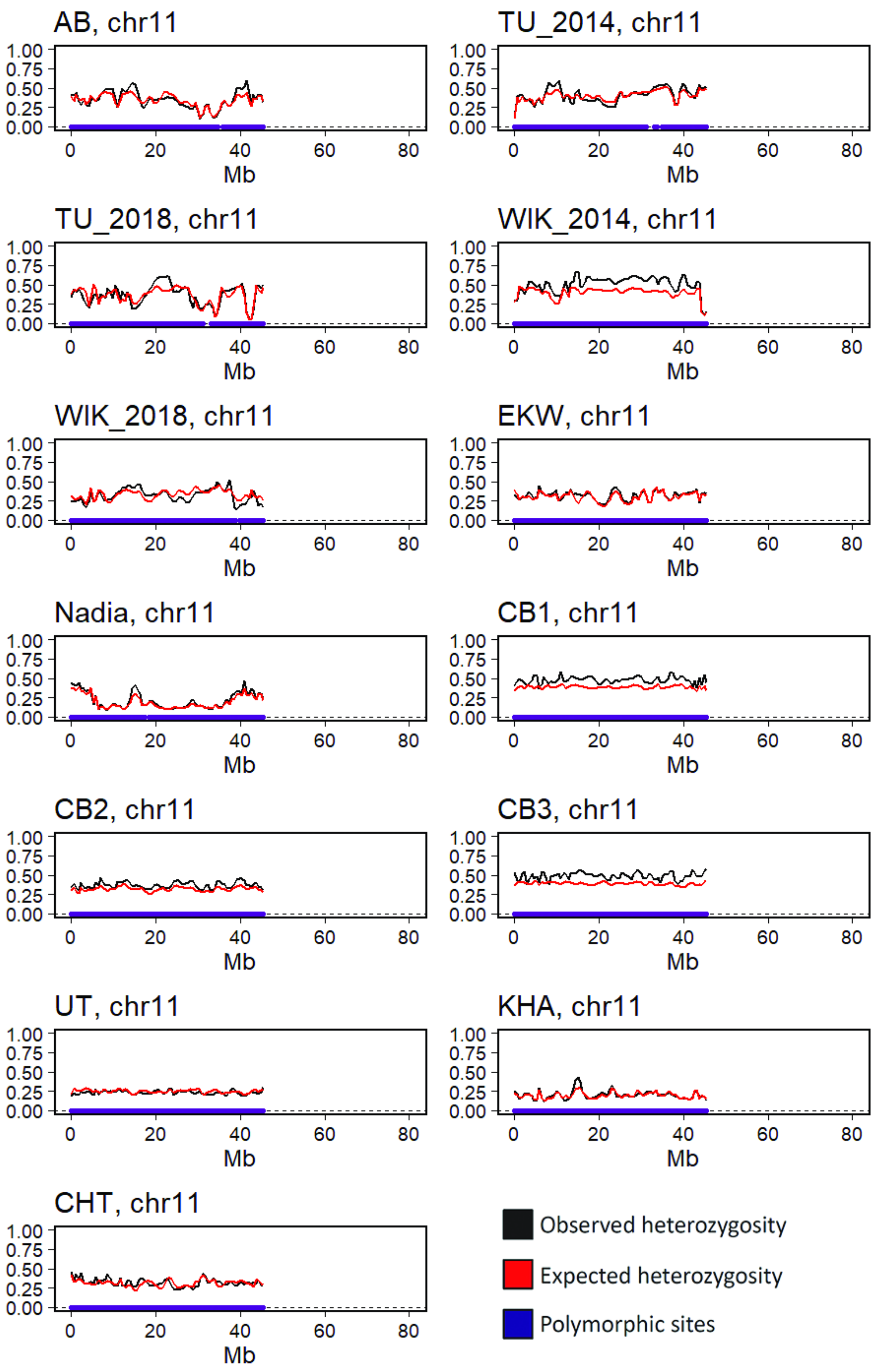
AB, chr11
TU_2014, chr11
TU_2018, chr11
WIK_2014, chr11
WIK_2018, chr11
EKW, chr11
Nadia, chr11
CB1, chr11
CB2, chr11
CB3, chr11
UT, chr11
KHA, chr11
CHT, chr11
1.00
0.75
0.50
0.25
0.00
0
20
40
60
80
Mb
Observed heterozygosity
Expected heterozygosity
Polymorphic sites

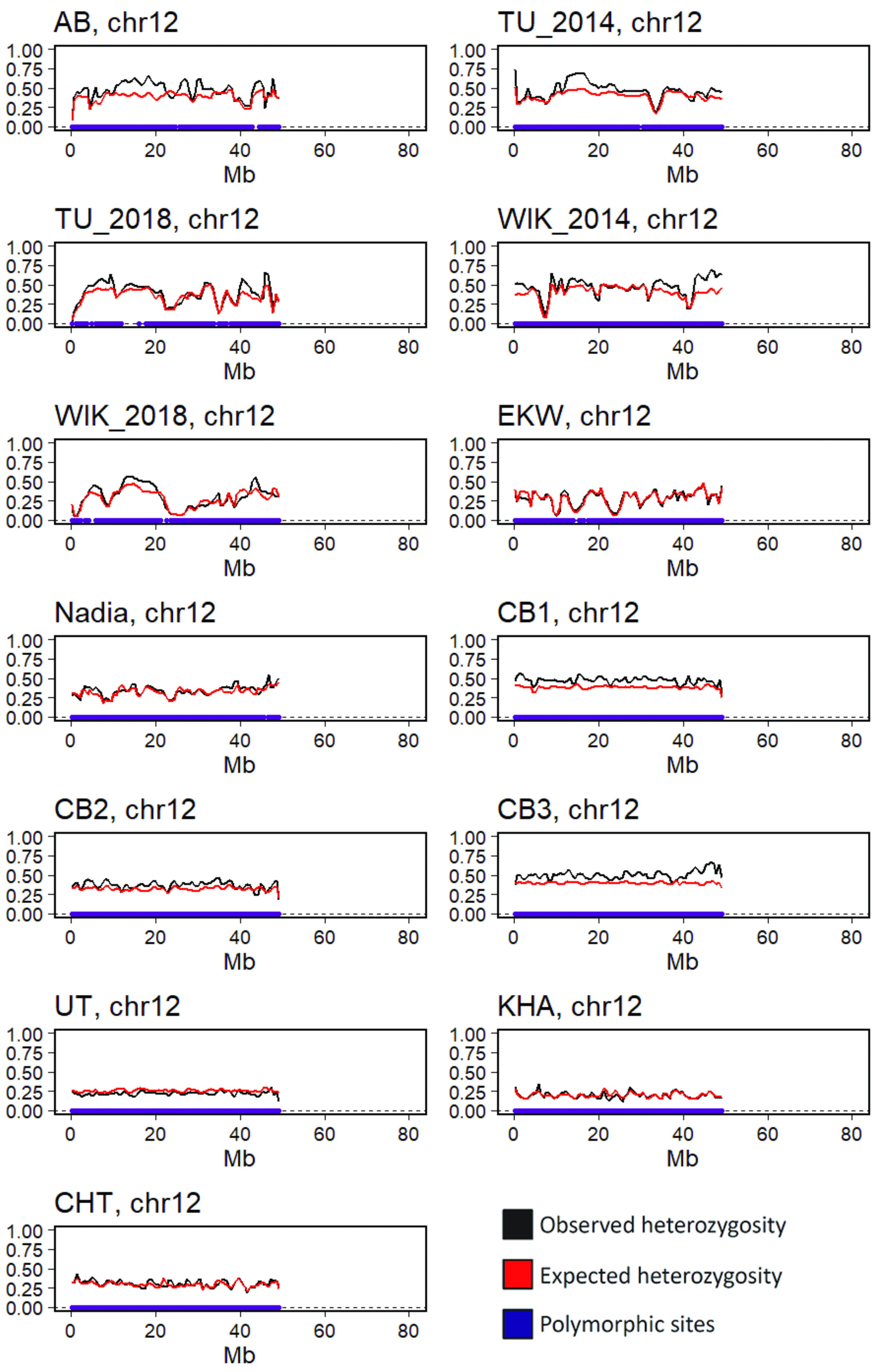
AB, chr12
TU_2014, chr12
TU_2018, chr12
WIK_2014, chr12
WIK_2018, chr12
EKW, chr12
Nadia, chr12
CB1, chr12
CB2, chr12
CB3, chr12
UT, chr12
KHA, chr12
CHT, chr12
0.00
0.25
0.50
0.75
1.00
0
20
40
60
80
Mb
Observed heterozygosity
Expected heterozygosity
Polymorphic sites
