## Supplemental Data S4 - heterozygosity chr13-25 for "The laboratory domestication of zebrafish: from diverse populations to inbred substrains"

Supplementary data S3/S4 for:

### The laboratory domestication of zebrafish: from diverse populations to inbred substrains

Jaanus Suurväli, Andrew R. Whiteley, Yichen Zheng, Karim Gharbi, Maria Leptin, Thomas Wiehe

Heterozygosity in wild and laboratory zebrafish.

Data was smoothed with Loess regression (span=0.06) prior to plotting it across chromosomes.

Supplementary Data S3: chromosomes 1-12

Supplementary Data S4: chromosomes 13-25

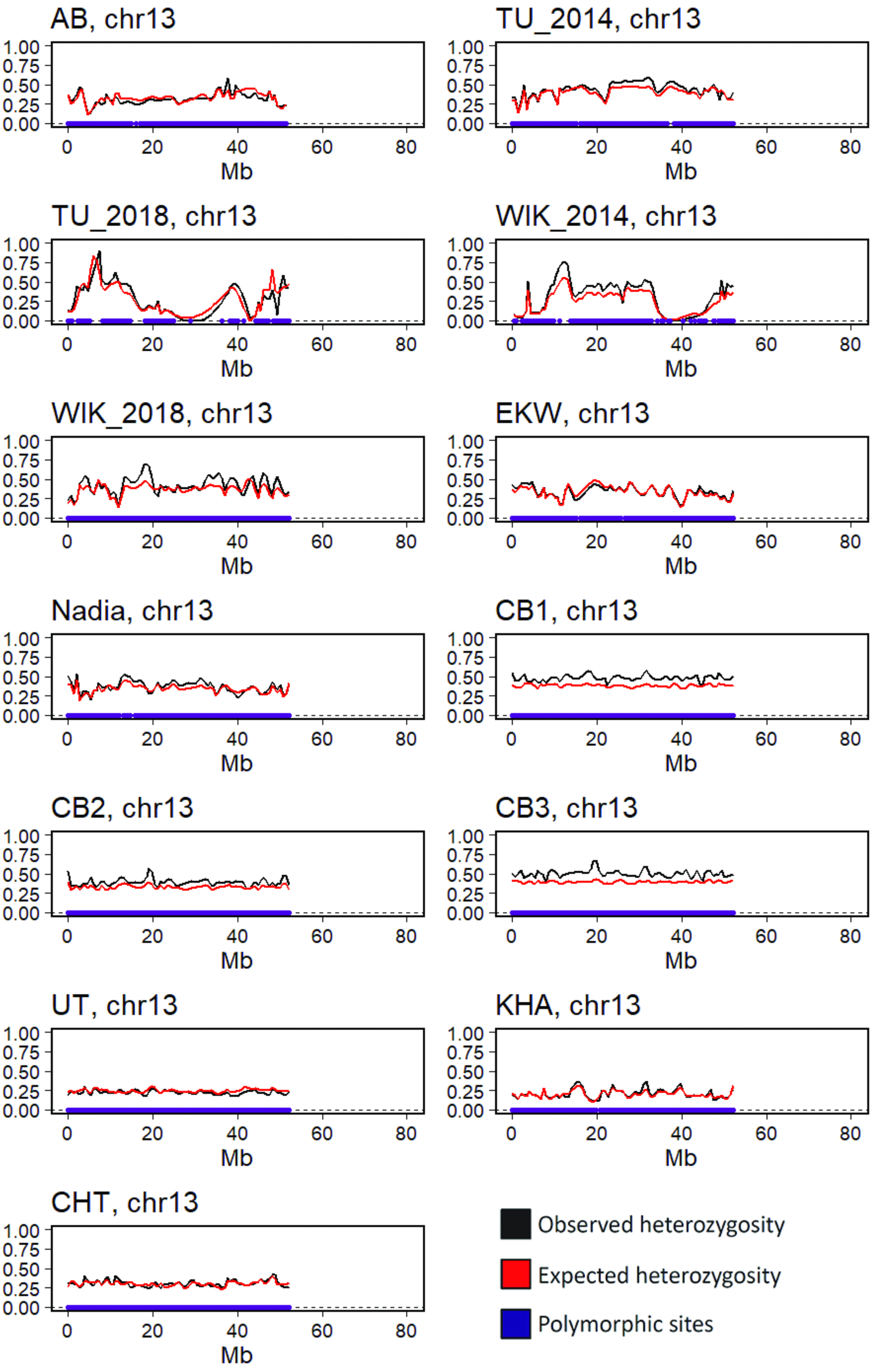

AB, chr13
TU_2014, chr13
TU_2018, chr13
WIK_2014, chr13
WIK_2018, chr13
EKW, chr13
Nadia, chr13
CB1, chr13
CB2, chr13
CB3, chr13
UT, chr13
KHA, chr13
CHT, chr13
1.00
0.75
0.50
0.25
0.00
0
20
40
60
80
Mb
Observed heterozygosity
Expected heterozygosity
Polymorphic sites

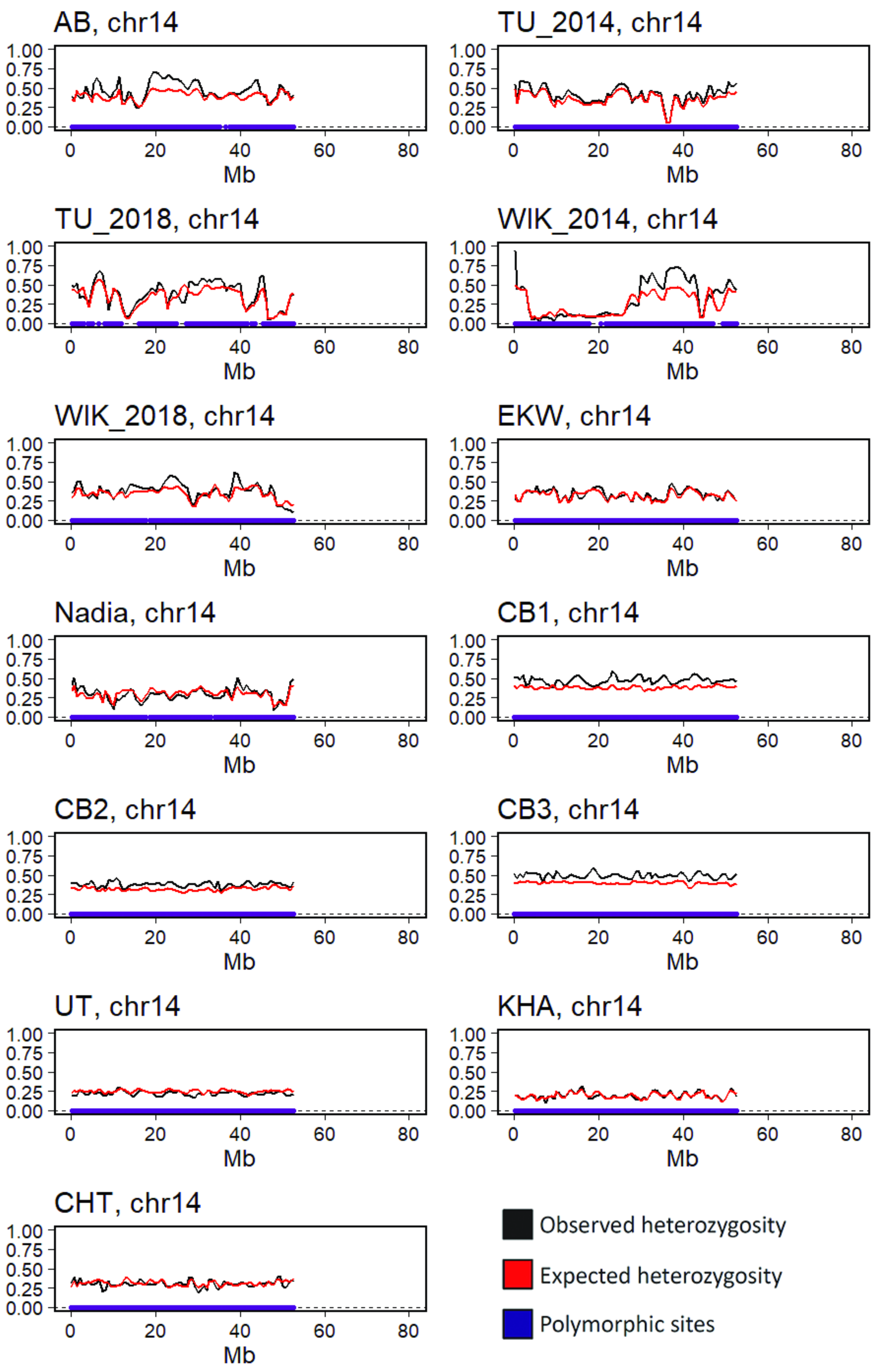
AB, chr14
TU_2014, chr14
TU_2018, chr14
WIK_2014, chr14
WIK_2018, chr14
EKW, chr14
Nadia, chr14
CB1, chr14
CB2, chr14
CB3, chr14
UT, chr14
KHA, chr14
CHT, chr14
1.00
0.75
0.50
0.25
0.00
0
20
40
60
80
Mb
Observed heterozygosity
Expected heterozygosity
Polymorphic sites

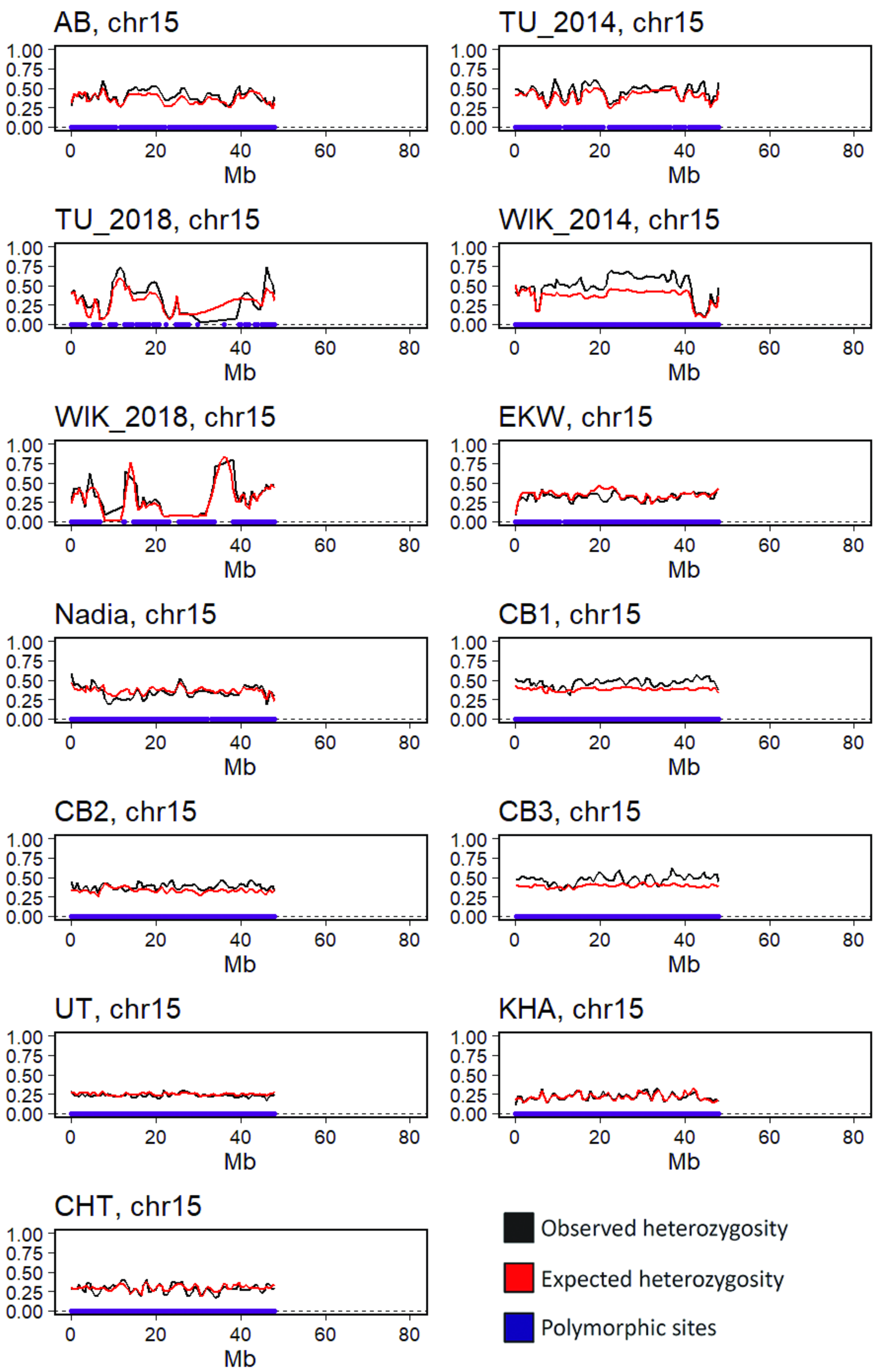
AB, chr15
TU_2014, chr15
TU_2018, chr15
WIK_2014, chr15
WIK_2018, chr15
EKW, chr15
Nadia, chr15
CB1, chr15
CB2, chr15
CB3, chr15
UT, chr15
KHA, chr15
CHT, chr15
1.00
0.75
0.50
0.25
0.00
0
20
40
60
80
Mb
Observed heterozygosity
Expected heterozygosity
Polymorphic sites

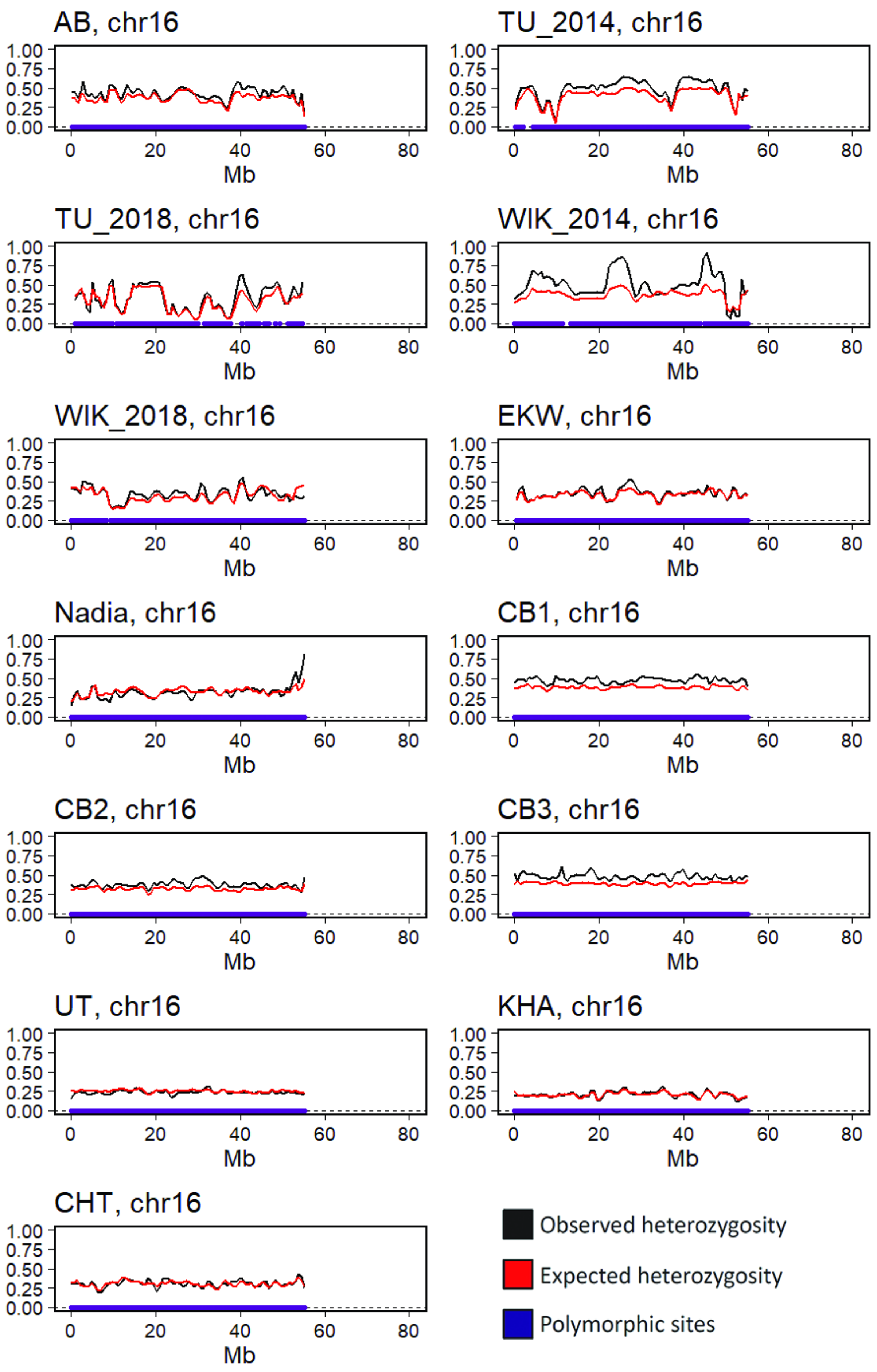
AB, chr16
TU_2014, chr16
TU_2018, chr16
WIK_2014, chr16
WIK_2018, chr16
EKW, chr16
Nadia, chr16
CB1, chr16
CB2, chr16
CB3, chr16
UT, chr16
KHA, chr16
CHT, chr16
1.00
0.75
0.50
0.25
0.00
0
20
40
60
80
Mb
Observed heterozygosity
Expected heterozygosity
Polymorphic sites

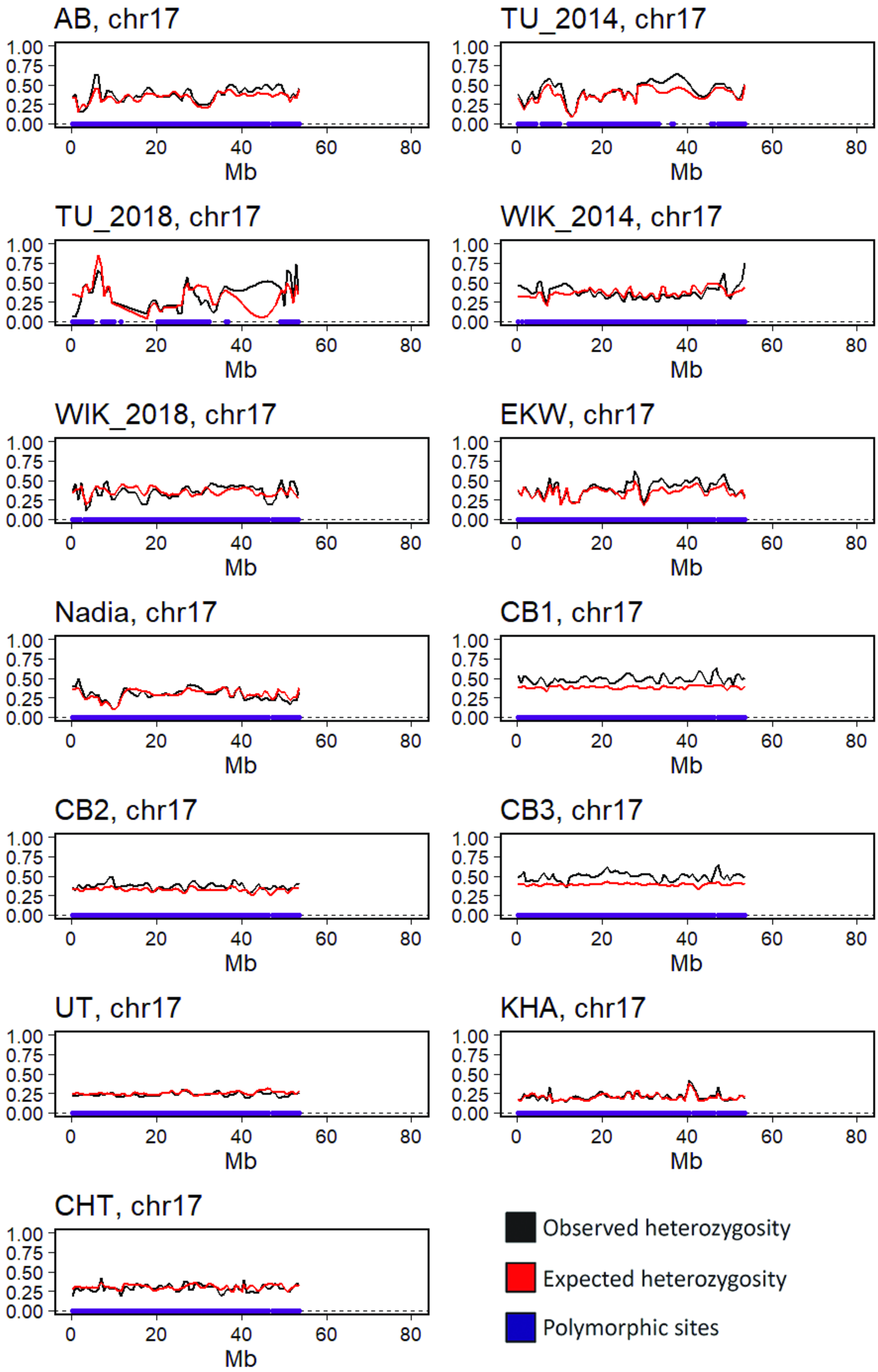

AB, chr17
TU_2014, chr17
TU_2018, chr17
WIK_2014, chr17
WIK_2018, chr17
EKW, chr17
Nadia, chr17
CB1, chr17
CB2, chr17
CB3, chr17
UT, chr17
KHA, chr17
CHT, chr17
1.00
0.75
0.50
0.25
0.00
0
20
40
60
80
Mb
Observed heterozygosity
Expected heterozygosity
Polymorphic sites

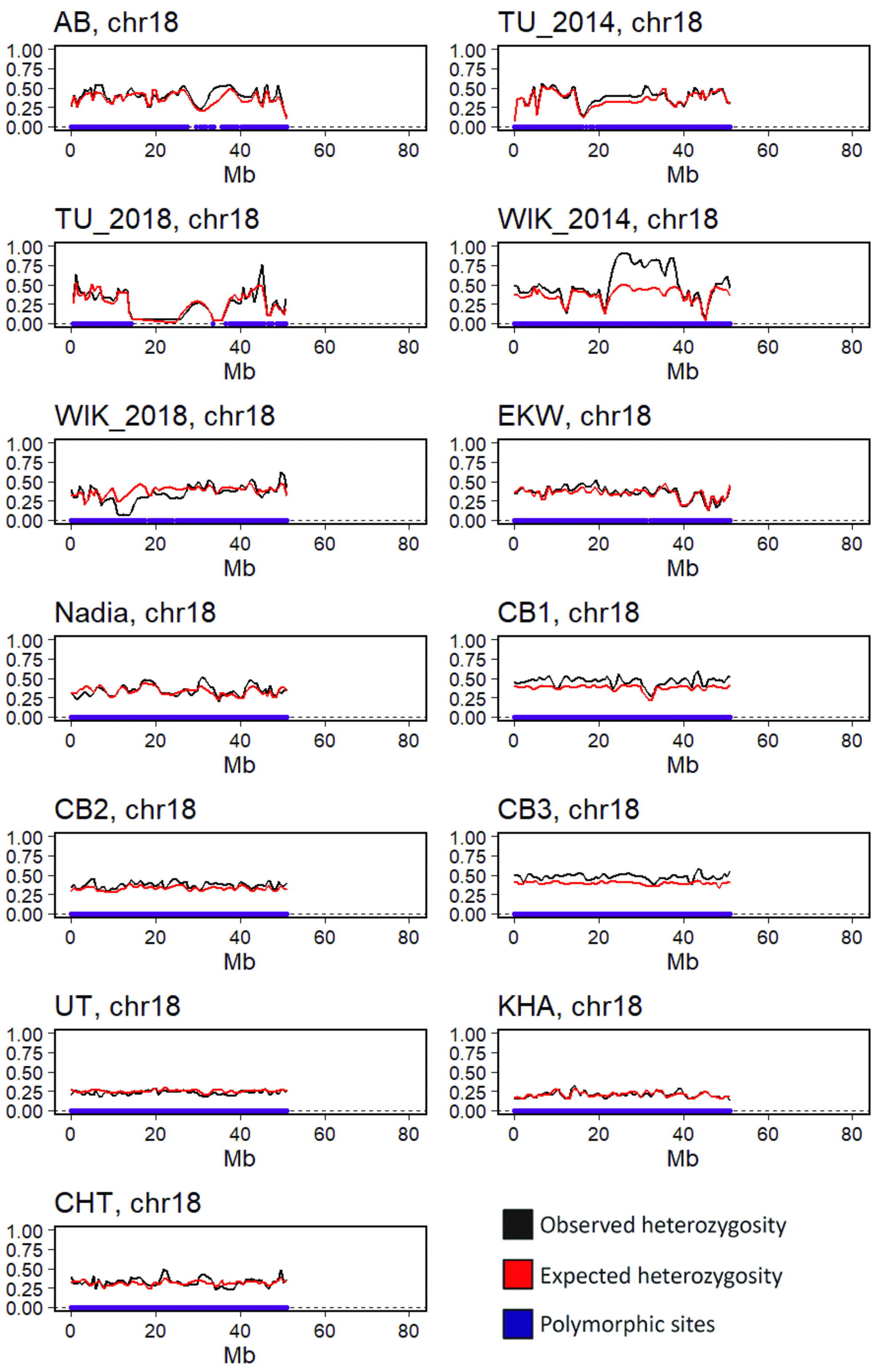

AB, chr18
TU_2014, chr18
TU_2018, chr18
WIK_2014, chr18
WIK_2018, chr18
EKW, chr18
Nadia, chr18
CB1, chr18
CB2, chr18
CB3, chr18
UT, chr18
KHA, chr18
CHT, chr18
1.00
0.75
0.50
0.25
0.00
0
20
40
60
80
Mb
Observed heterozygosity
Expected heterozygosity
Polymorphic sites

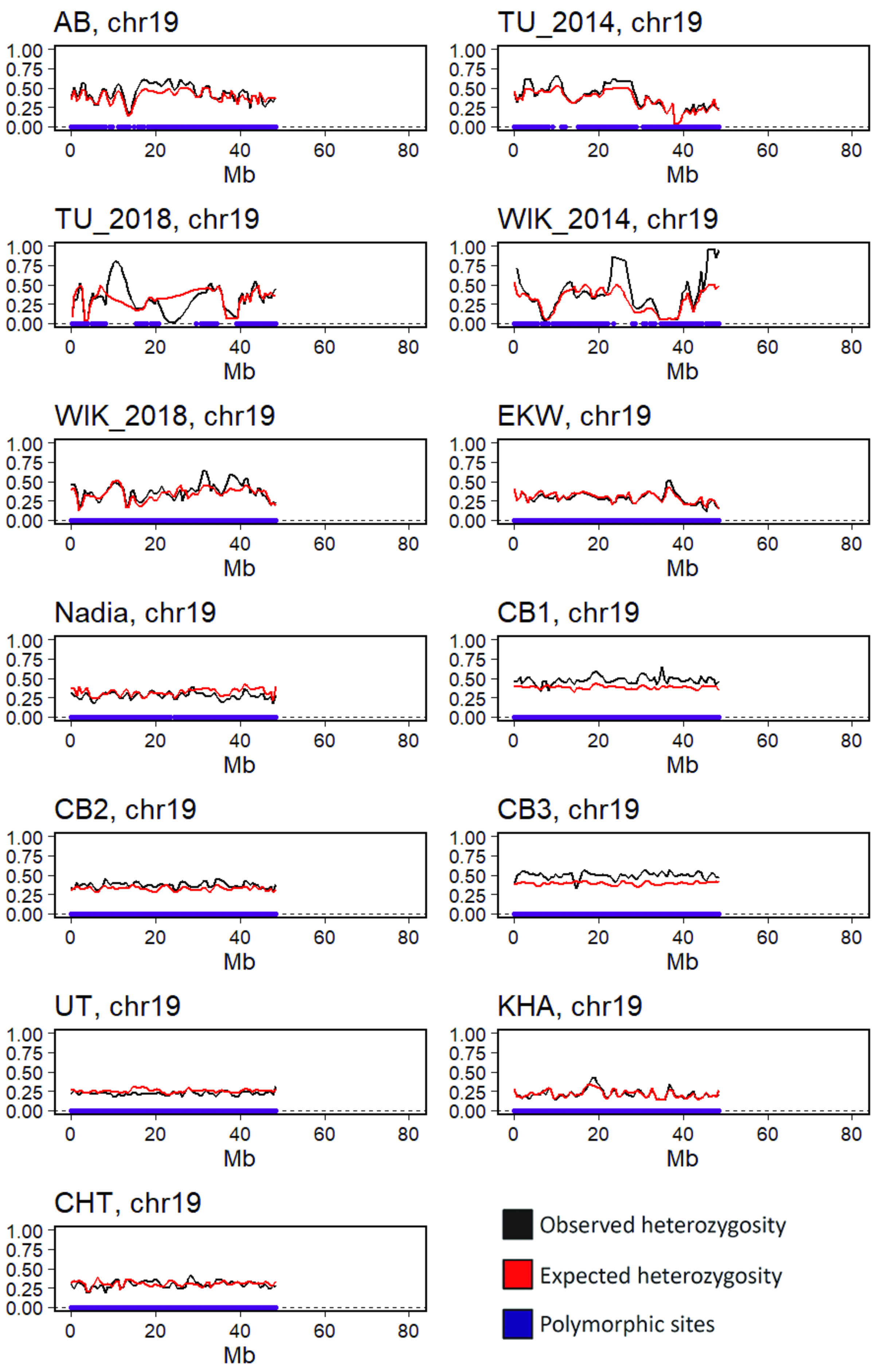
AB, chr19
TU_2014, chr19
TU_2018, chr19
WIK_2014, chr19
WIK_2018, chr19
EKW, chr19
Nadia, chr19
CB1, chr19
CB2, chr19
CB3, chr19
UT, chr19
KHA, chr19
CHT, chr19
1.00
0.75
0.50
0.25
0.00
0
20
40
60
80
Mb
Observed heterozygosity
Expected heterozygosity
Polymorphic sites

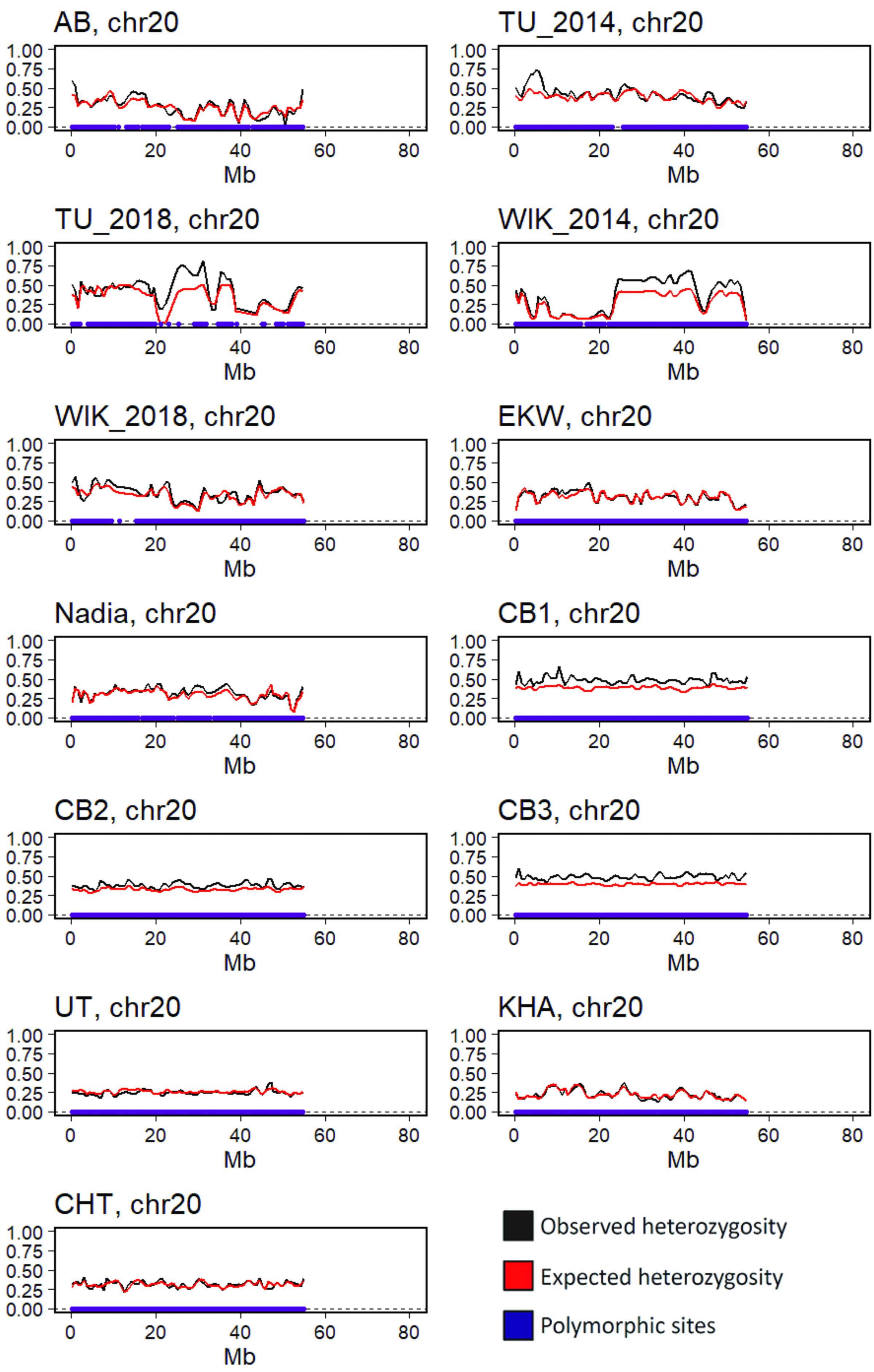
AB, chr20
TU_2014, chr20
TU_2018, chr20
WIK_2014, chr20
WIK_2018, chr20
EKW, chr20
Nadia, chr20
CB1, chr20
CB2, chr20
CB3, chr20
UT, chr20
KHA, chr20
CHT, chr20
1.00
0.75
0.50
0.25
0.00
0
20
40
60
80
Mb
Observed heterozygosity
Expected heterozygosity
Polymorphic sites

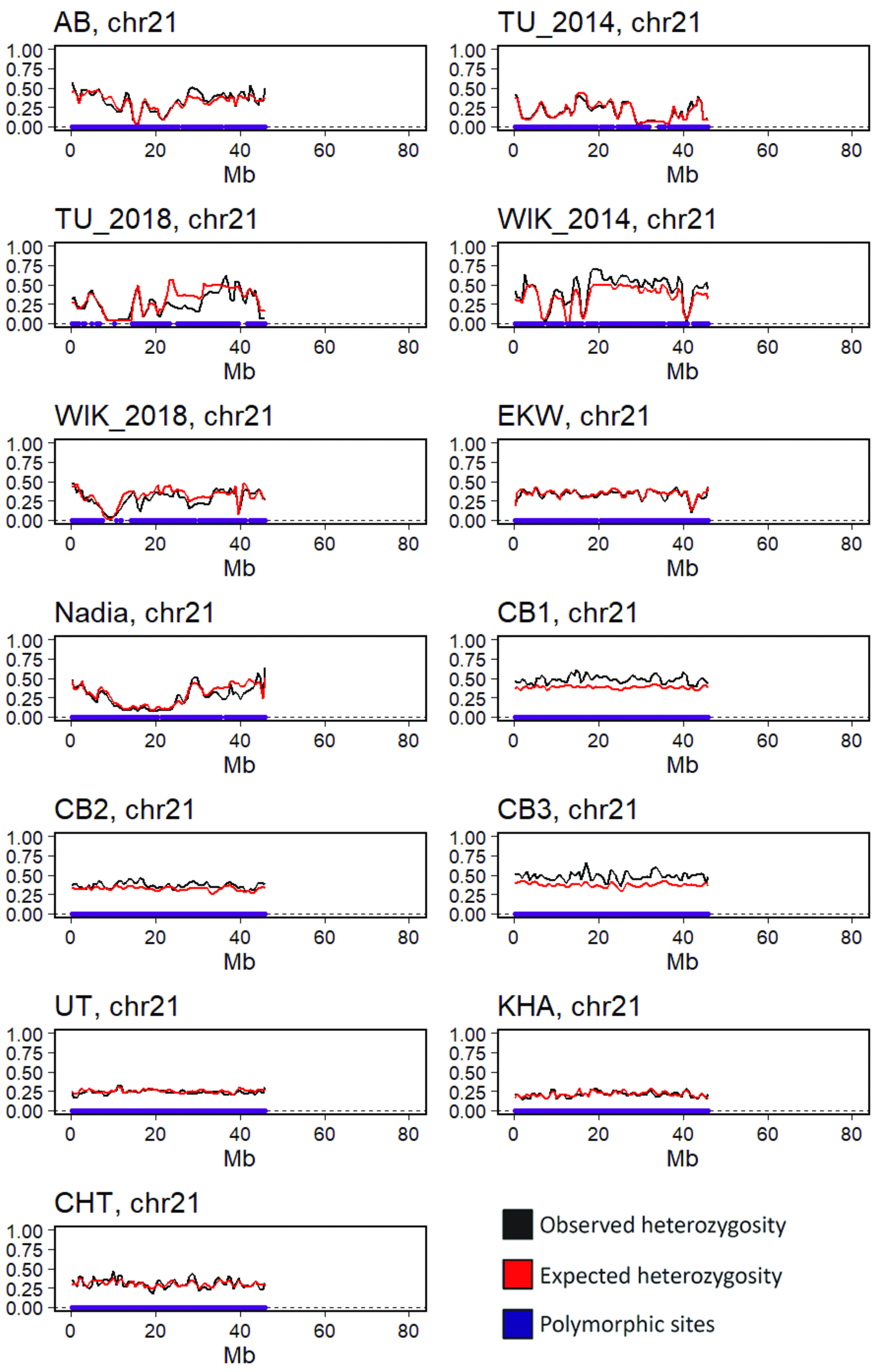
AB, chr21
TU_2014, chr21
TU_2018, chr21
WIK_2014, chr21
WIK_2018, chr21
EKW, chr21
Nadia, chr21
CB1, chr21
CB2, chr21
CB3, chr21
UT, chr21
KHA, chr21
CHT, chr21
1.00
0.75
0.50
0.25
0.00
0
20
40
60
80
Mb
Observed heterozygosity
Expected heterozygosity
Polymorphic sites

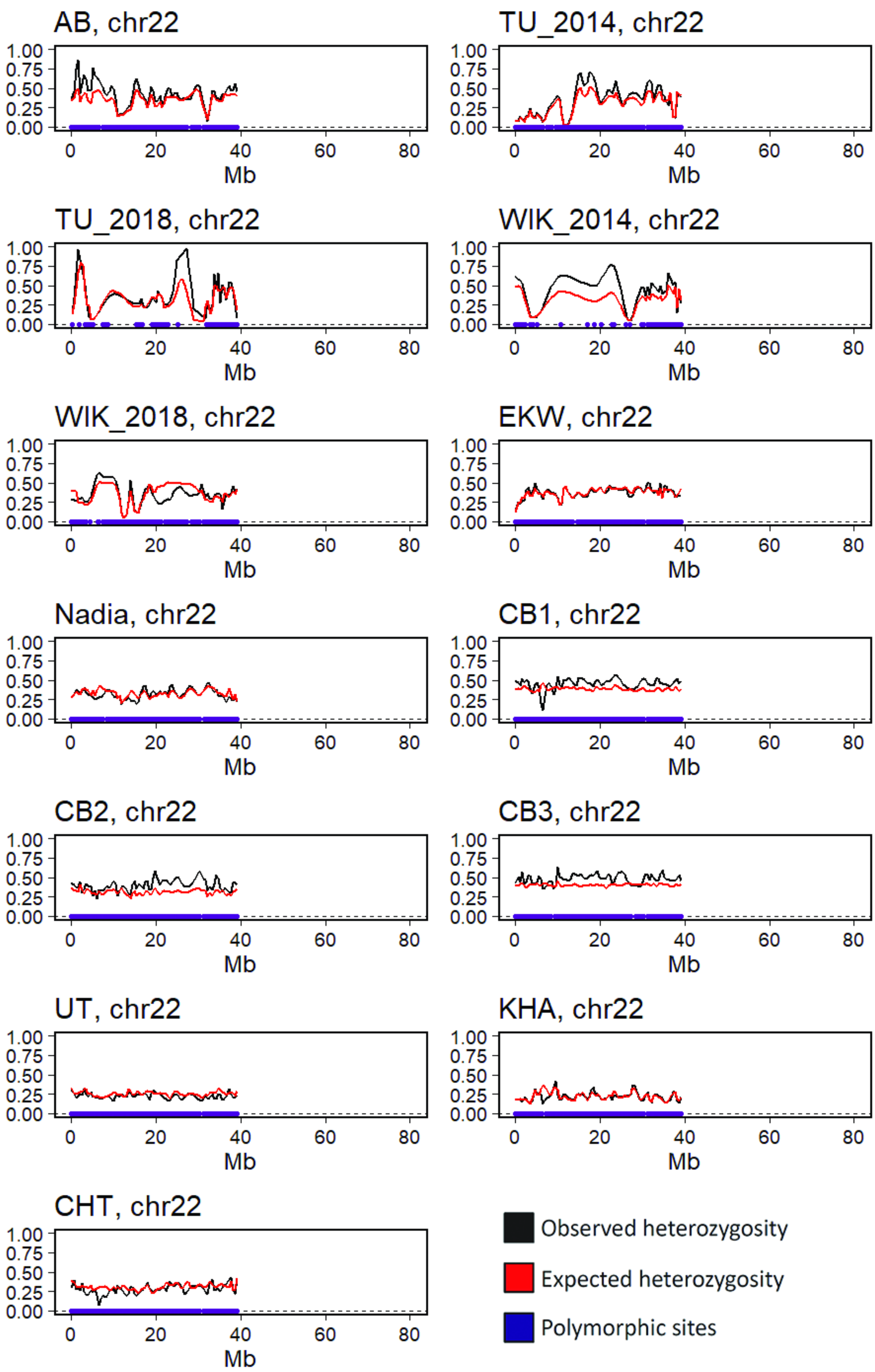

AB, chr22
TU_2014, chr22
TU_2018, chr22
WIK_2014, chr22
WIK_2018, chr22
EKW, chr22
Nadia, chr22
CB1, chr22
CB2, chr22
CB3, chr22
UT, chr22
KHA, chr22
CHT, chr22
Mb
Observed heterozygosity
Expected heterozygosity
Polymorphic sites

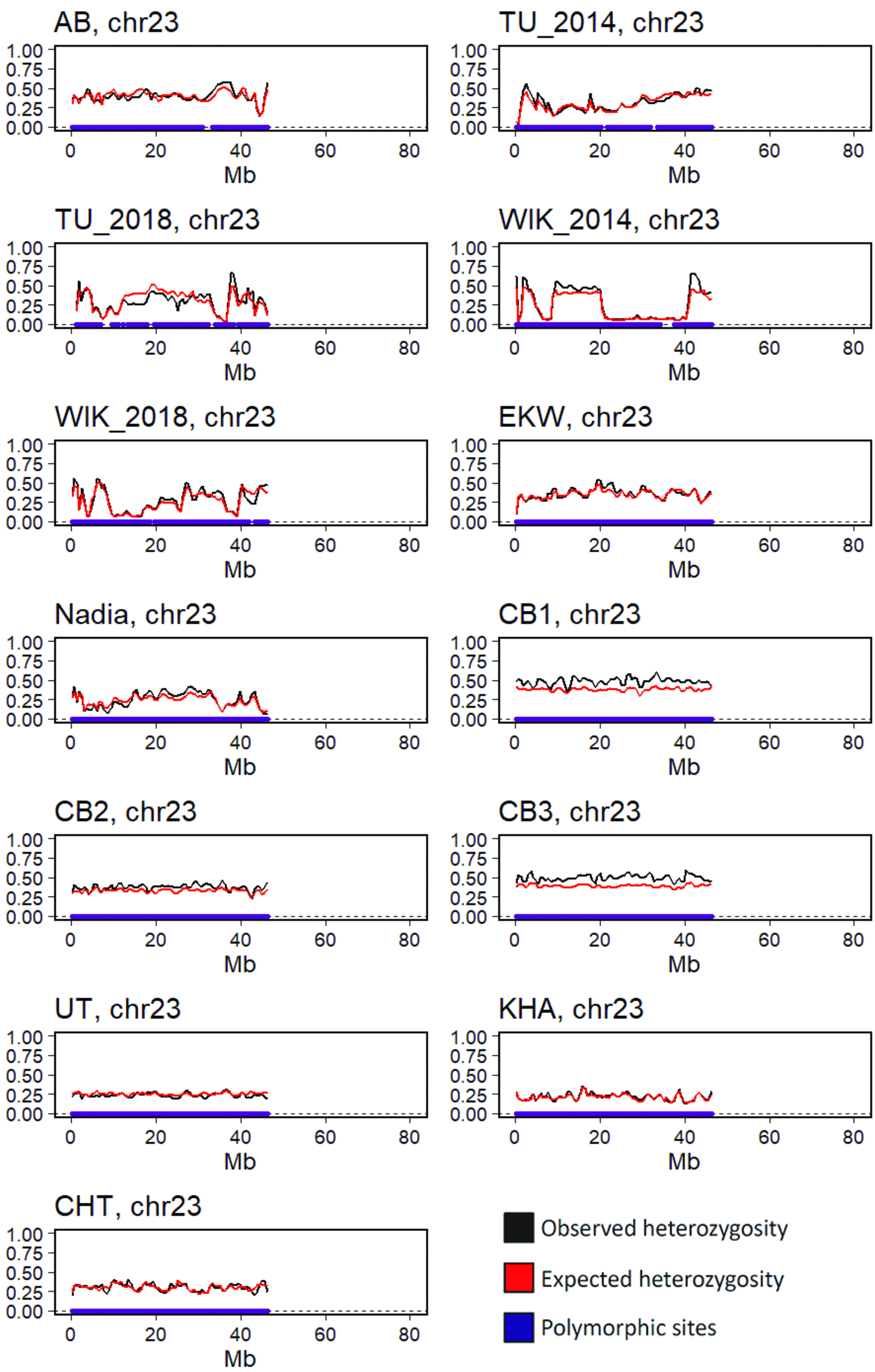

AB, chr23
TU_2014, chr23
TU_2018, chr23
WIK_2014, chr23
WIK_2018, chr23
EKW, chr23
Nadia, chr23
CB1, chr23
CB2, chr23
CB3, chr23
UT, chr23
KHA, chr23
CHT, chr23
1.00
0.75
0.50
0.25
0.00
0
20
40
60
80
Mb
Observed heterozygosity
Expected heterozygosity
Polymorphic sites

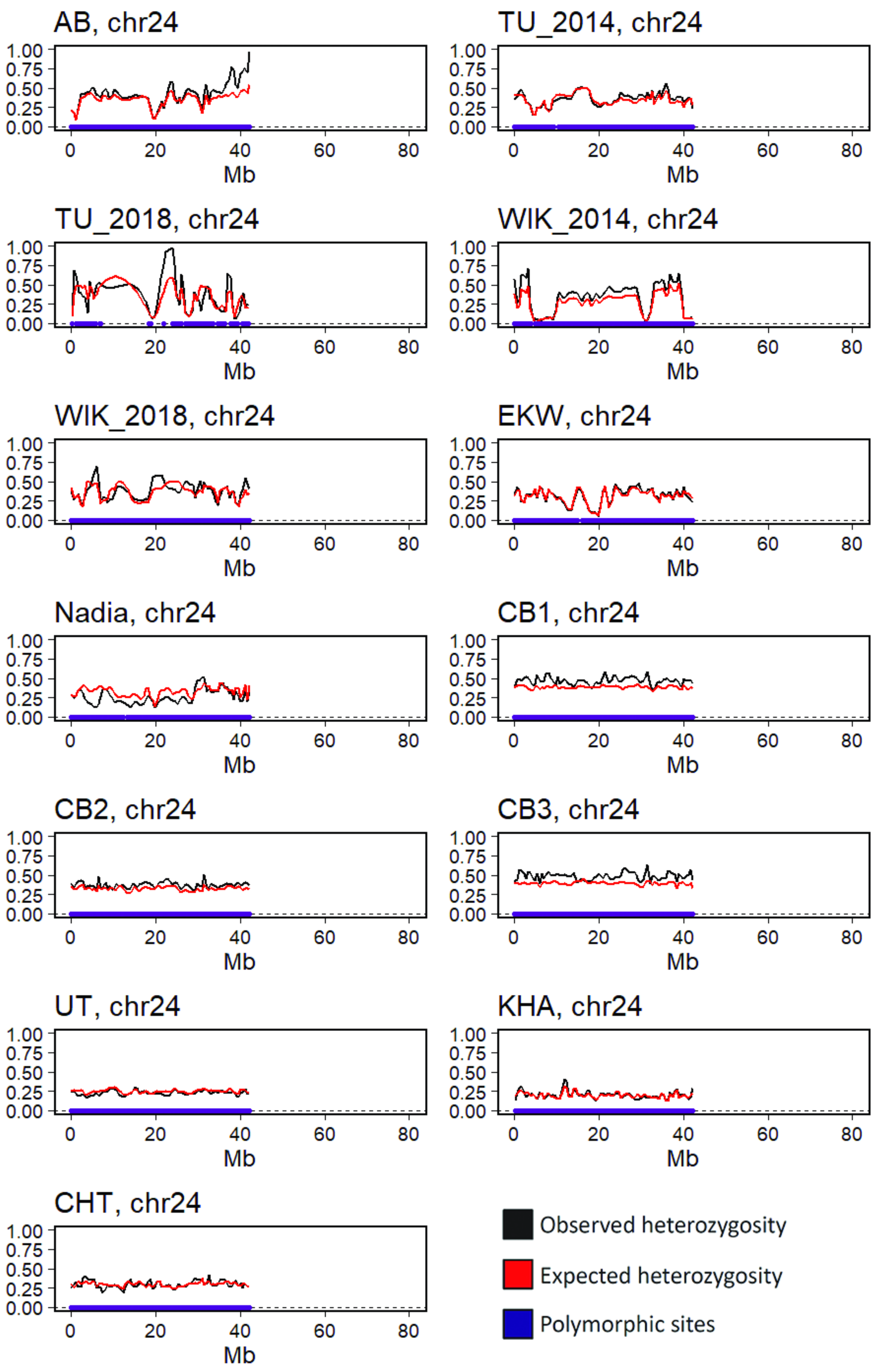
AB, chr24
TU_2014, chr24
TU_2018, chr24
WIK_2014, chr24
WIK_2018, chr24
EKW, chr24
Nadia, chr24
CB1, chr24
CB2, chr24
CB3, chr24
UT, chr24
KHA, chr24
CHT, chr24
1.00
0.75
0.50
0.25
0.00
0
20
40
60
80
Mb
Observed heterozygosity
Expected heterozygosity
Polymorphic sites

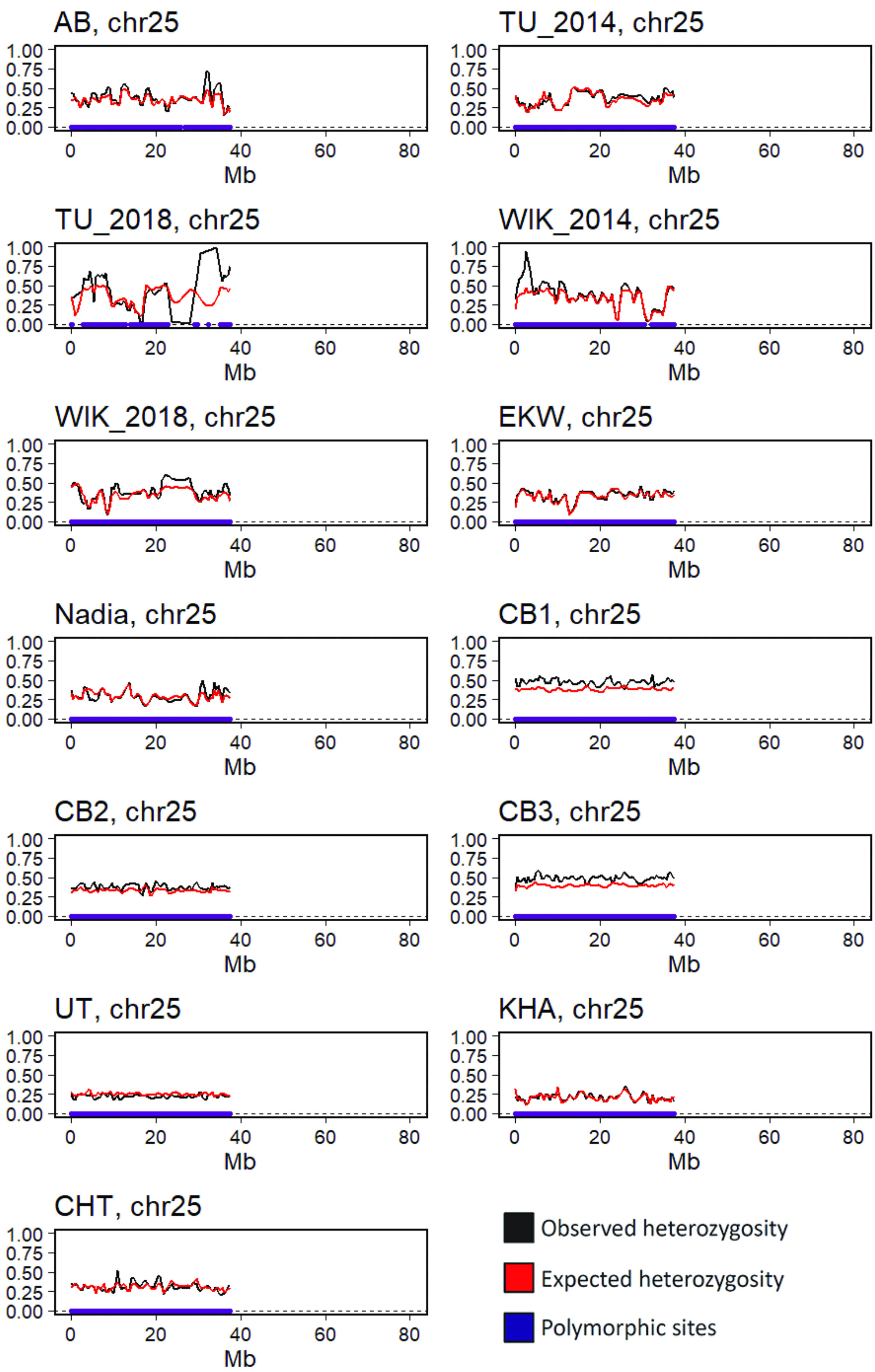

AB, chr25
TU_2014, chr25
TU_2018, chr25
WIK_2014, chr25
WIK_2018, chr25
EKW, chr25
Nadia, chr25
CB1, chr25
CB2, chr25
CB3, chr25
UT, chr25
KHA, chr25
CHT, chr25
1.00
0.75
0.50
0.25
0.00
0
20
40
60
80
Mb
Observed heterozygosity
Expected heterozygosity
Polymorphic sites
