## Supplemental Data S5 - frequency spectra for "The laboratory domestication of zebrafish: from diverse populations to inbred substrains"

Supplementary data S5 for:

### The laboratory domestication of zebrafish: from diverse populations to inbred substrains

Jaanus Suurväli, Andrew R. Whiteley, Yichen Zheng, Karim Gharbi, Maria Leptin, Thomas Wiehe

Allele frequency (AF) spectra in wild and laboratory zebrafish.

(1) Spectra for the derived AF, with the most commonly observed allele in the dataset as reference

(2) Spectra for the folded AF, e.g. the least frequent allele in a given population.

(3) Spectra for the derived AF, with mutations that increase or decrease GC content shown separately. The reference allele was defined here as "most commonly observed allele in a dataset of 7 fish each from UT, KHA and CHT"

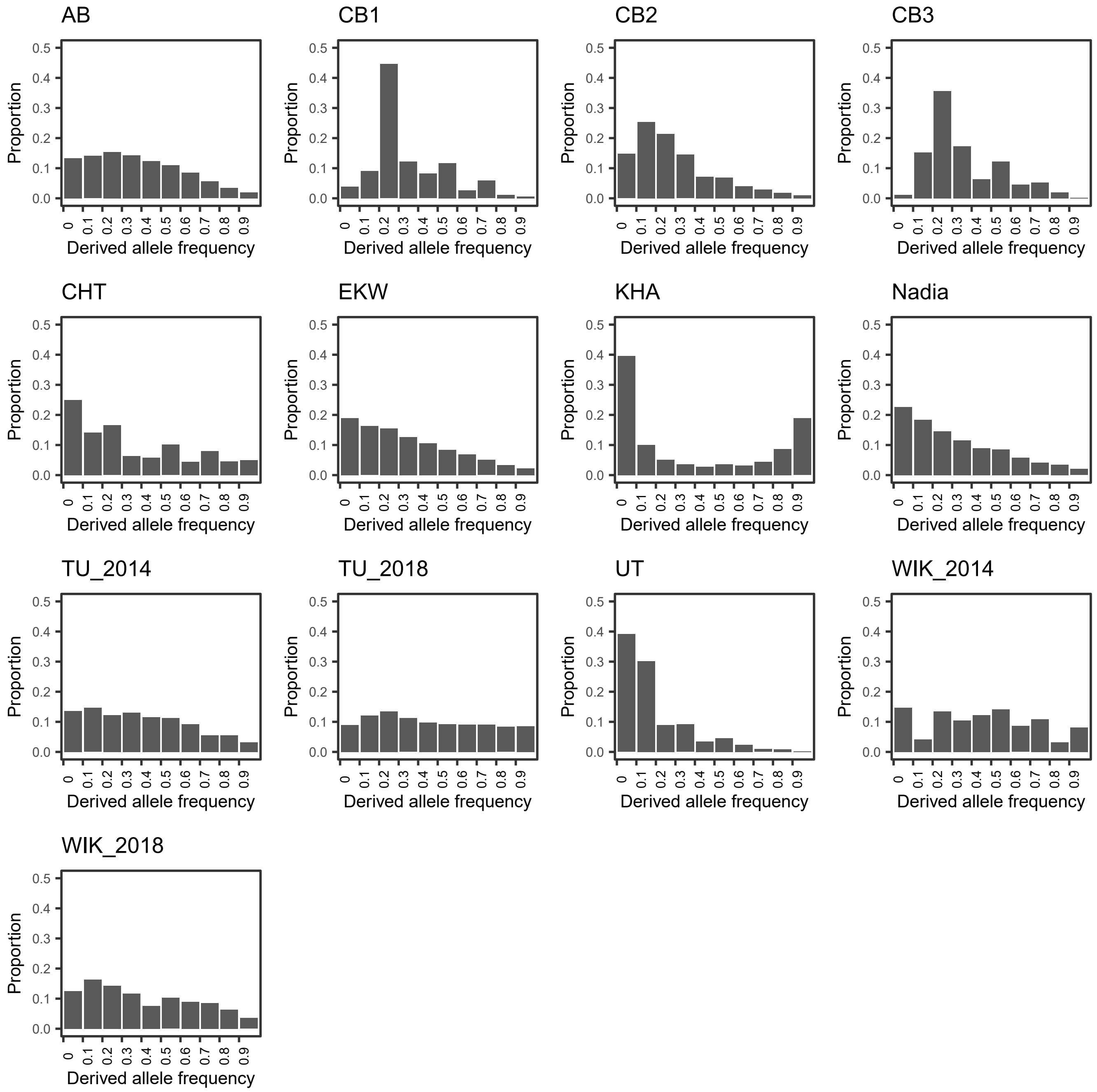


Derived allele frequency spectra for wild and laboratory zebrafish.

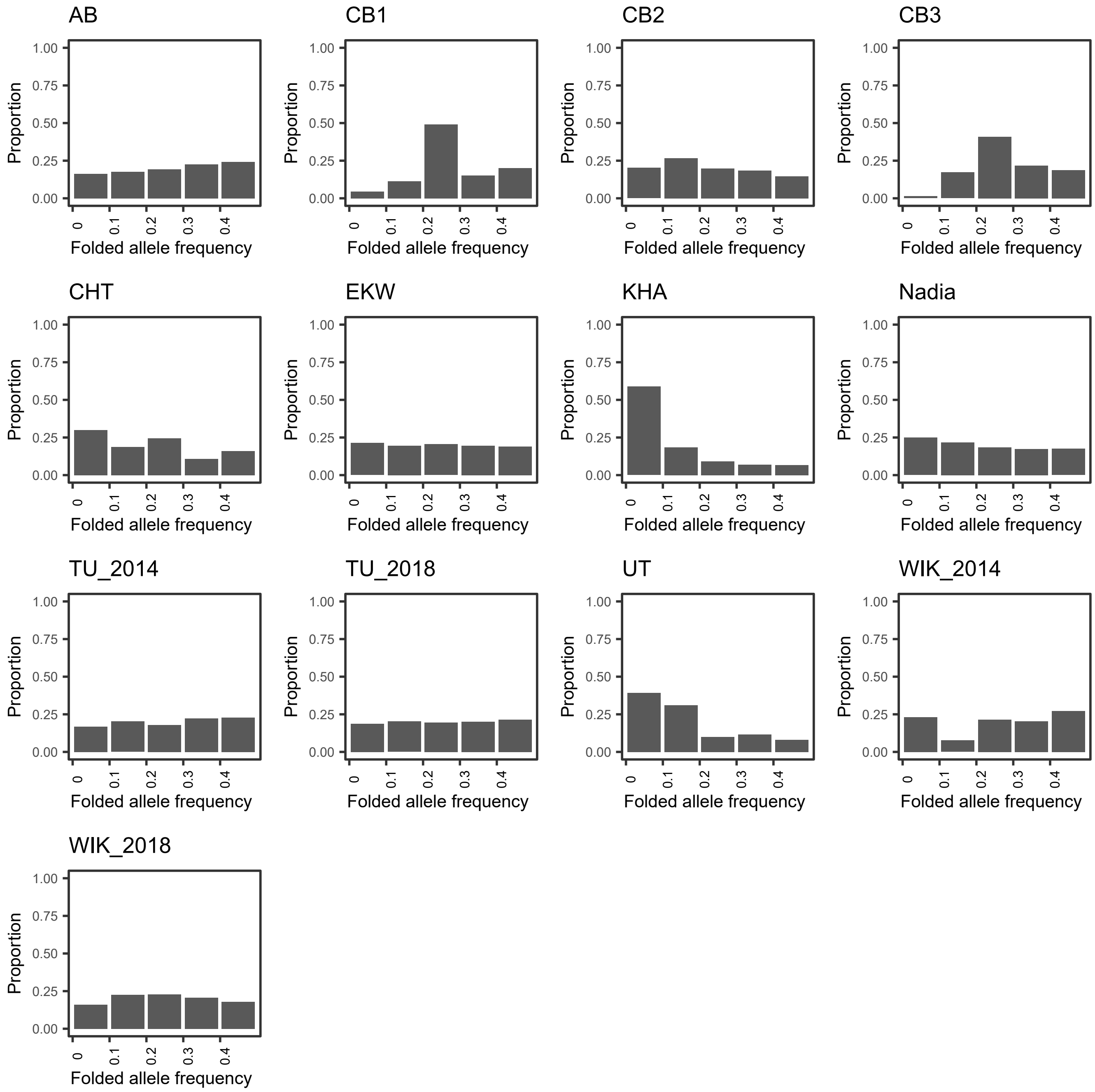


Folded allele frequency spectra for wild and laboratory zebrafish.

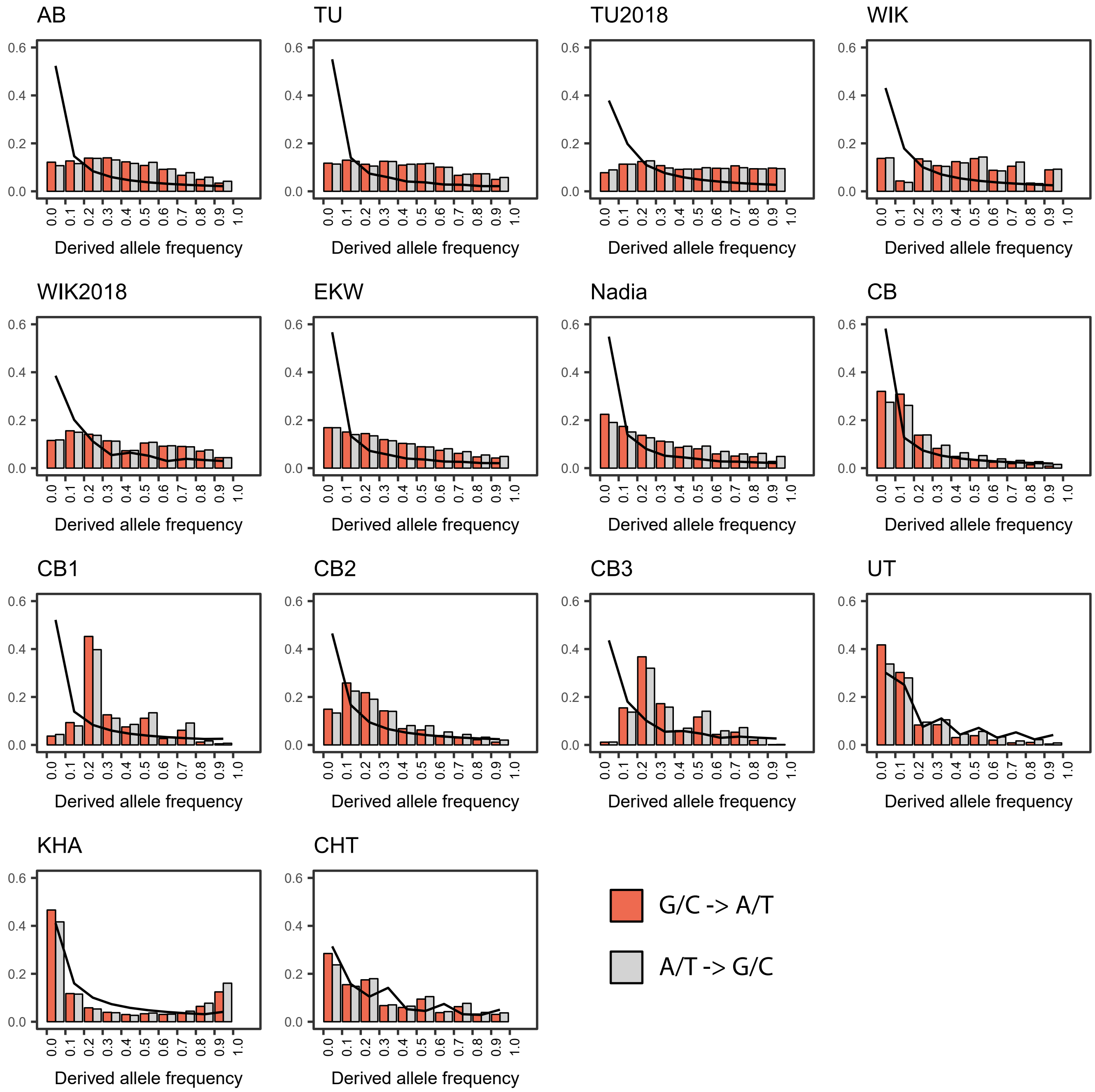


Derived frequency spectra, calculated with reference allele defined as the most commonly observed allele in 7 individuals from each of the three wild populations.
